# Arousal dependent modulation of thalamo-cortical functional interaction

**DOI:** 10.1101/135376

**Authors:** Iain Stitt, Zhe Charles Zhou, Susanne Radtke-Schuller, Flavio Fröhlich

## Abstract

Cognition and behavior emerge from the dynamic interaction of widely distributed, but functionally specialized brain networks. However, it remains unclear how network-level interactions dynamically reorganize to support ever-shifting cognitive and behavioral demands. Here, we investigate how the interaction between posterior parietal cortex (PPC) and lateral posterior (LP) / Pulvinar is shaped by ongoing fluctuations in pupil-linked arousal, which is a non-invasive measure related to neuromodulatory tone in the brain. We found that fluctuations in pupil-linked arousal tracked the dynamic interaction between PPC and LP/Pulvinar characterized by changes in the direction and carrier frequency of oscillatory interaction. Active visual exploration by saccadic eye movements elicited similar transitions in thalamo-cortical interaction. These findings suggest a common network substrate of both spontaneous activity and active vision. Thus, neuromodulators may play a role in dynamically sculpting the patterns of thalamo-cortical functional interaction that underlie visual processing.

## Introduction

As exemplified by the daydreaming student drifting in and out of focus during class, the brain exhibits the ability to rapidly transition between states of varying engagement with the external world. Rather than resulting from changes in anatomical connections between neurons, such moment-to-moment variability in internal brain state arises through changes in network-level activity patterns occurring within the brains structural framework (Deco et al., 2011). In this scheme, cognition and behavior emerge from the dynamic interaction of widely distributed, but functionally specialized cortical and subcortical brain regions (Koch et al., 2016; Siegel et al., 2012).

In the visual system, higher order brain structures such as the lateral posterior (LP) / Pulvinar nuclear complex of the thalamus play an essential role in orchestrating the patterns of large-scale cortical interaction that underlie visual behavior (Jones, 2001; Saalmann and Kastner, 2011; Saalmann et al., 2012). Providing the structural framework for this coordinating role, LP/Pulvinar exhibits reciprocal anatomical connectivity with widely distributed visual cortical areas (Jones, 2001). In particular, poster parietal cortex (PPC), a higher-order association area that itself is a hub of sensory integration, sends dense projections to PL/Pulvinar (Manger et al., 2002). Converging evidence suggests that the selective synchronization of neuronal oscillations between LP/Pulvinar and cortex facilitates communication between inter-connected cortical sites, and that such patterns of thalamo-cortical dynamics reflect the circuit-level computations that underlie visual sensory processing and behavior (Fries, 2015; Siegel et al., 2012). Yet, despite the established importance of thalamo-cortical interaction for visual processing and behavior (Saalmann and Kastner, 2011; Saalmann et al., 2012), we still lack a clear understanding of how the relative strength of thalamus-to-cortex and cortex-to-thalamus signals are dynamically tuned to meet current behavioral demands.

Recent theoretical work proposed neuromodulators as a mechanism for modifying information flow in neuronal networks on a short temporal scale (Heeger, 2017). Indeed, the neuromodulators noradrenaline and acetylcholine modulate the intrinsic properties of neurons across both cortical and thalamic areas (McCormick, 1989; McCormick et al., 1993; Pape and McCormick, 1989; Polack et al., 2013; Rogawski and Aghajanian, 1980; Zagha and McCormick, 2014). Beyond the effects of such neurotransmitters on the local cellular level in cortex and thalamus, it remains unclear if neuromodulators help to shape emergent patterns of information routing in thalamo-cortical networks. Recent findings have linked ongoing fluctuations in pupil diameter to the release of noradrenaline and acetylcholine from synaptic terminals in the cortex (Reimer et al., 2016), indicating that non-invasive measurement of pupil-linked arousal enables the indirect inference of neuromodulatory tone in the brain.

To investigate the role of neuromodulation in shaping thalamo-cortical functional interaction, we monitored ongoing fluctuations in pupil-linked arousal in awake head-restrained ferrets, while simultaneously recording spiking and local field potential (LFP) activity from LP/Pulvinar and PPC (Fig. 1a). We found that the carrier frequency of thalamo-cortical synchronization varied with arousal, and that the direction of causal interaction between cortex and thalamus switched between low and high arousal levels. Moreover, we show that such transitions in network dynamics are not exclusive to ongoing activity in relative absence of visual input, but also occur when animals are actively engaged in sensory processing. We suggest that neuromodulators shape thalamo-cortical functional interaction by altering the relative contribution of thalamic and cortical signals to emergent network dynamics.

**Figure 1 /.**
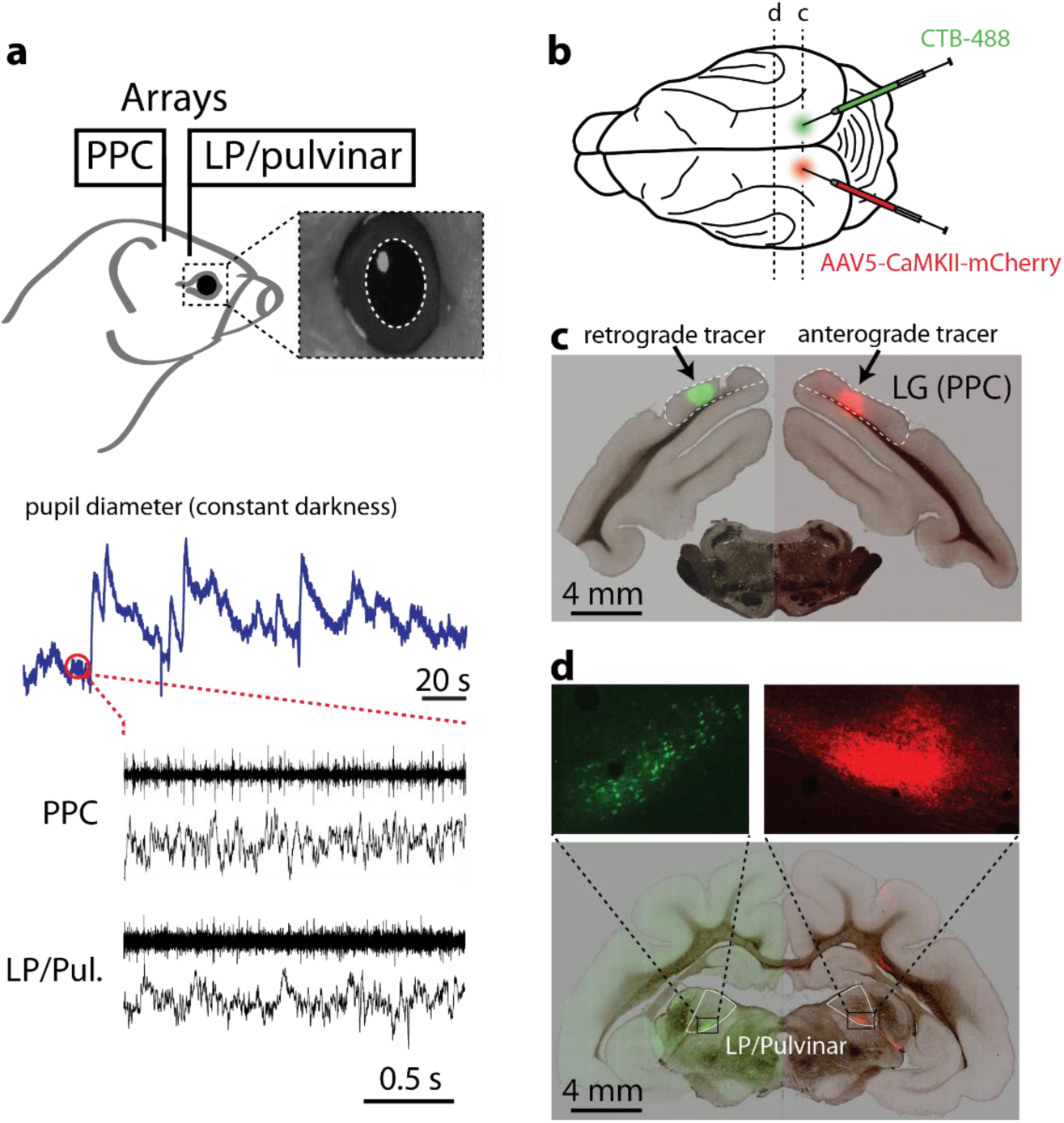
Experimental setup and anatomical connectivity between LP/Pulvinar and PPC. **a,** Diagram illustrating how neural signals from PPC and LP/Pulvinar were simultaneously recorded with pupil diameter. Inset image shows a typical view of the ferret infra-red eye tracking, with the pupil outlined in white. Below are raw traces of ongoing fluctuations pupil diameter and co-recorded spiking and LFP activity in PPC and LP/Pulvinar. Note that pupil diameter spontaneously fluctuates on both short and long time scales. **b**, Anterograde (rAAV5-CaMKII-mCherry) and retrograde (CTB-488) tracers were injected into PPC in the left and right hemispheres, respectively. **c**, Brightfield image of a brain section containing the PPC injection sites overlaid with green and red fluorescence channels. Fluorescent blobs show the location of anterograde and retrograde tracer in PPC. **d**, Brightfield image of thalamus overlaid with fluorescence from red and green channels. Retrograde labeling of cell bodies (green) and anterograde labeling of axonal projections (red) in corresponding locations of LP/Pulvinar illustrate reciprocal connectivity between PPC and LP/Pulvinar in the ferret. Abbreviations: LG lateral gyrus; PPC posterior parietal cortex.

## Results

### Reciprocal anatomical connectivity between PPC and LP/Pulvinar in the ferret

Functional interaction between brain regions is constrained by structural connectivity. To map the precise anatomical connectivity of the regions of interest in this study, we injected anterograde (AAV5-CaMKII-mCherry) and retrograde (CTB-488) tracers into the left and right PPC, respectively (Fig. 1b,c). PPC injections were made at locations that corresponded to the site of cortical multielectrode array implantation in other animals (Supplementary Fig. 1). We observed anterograde labeled fibers in the ventral portion of LP/Pulvinar (Fig. 1d) indicating that PPC neurons send projections to this sub region of the thalamus. In addition, we observed retrograde labeled cell bodies at the corresponding location in the opposite hemisphere (Fig. 1d), indicating that LP/Pulvinar also projects back to the location of the injection site in PPC. These results indicate that regions of thalamus and cortex where electrophysiological recordings were obtained display reciprocal connectivity, establishing the physical substrate for studying how thalamo-cortical interaction varies with pupil-linked arousal.

### Pupil-linked arousal dependent changes in neuronal spiking and LFP spectra

Given that both PPC and LP/Pulvinar receive dense projections from brainstem neuromodulatory systems (Foote and Morrison, 1987), we first examined how firing rate was modulated with pupil-linked arousal. Large and small pupil sizes indicated high and low arousal states, respectively. Consistent with previous *in vitro* and *in vivo* work on the influence of noradrenaline on neuronal excitability (McCormick, 1989; McCormick et al., 1993; McGinley et al., 2015a; McGinley et al., 2015b; Pape and McCormick, 1989; Polack et al., 2013; Rogawski and Aghajanian, 1980; Salgado et al., 2016), we found that neuronal firing rate at the population level in both PPC and LP/Pulvinar significantly increased with pupil diameter (Fig. 2b population, PPC: P = 0.003 one-way ANOVA, r = 0.30, LP/Pulvinar: P = 9.56 × 10^-11^, r = 0.55). Multi-unit spiking activity in PPC could be more generally separated into three groups based on correlation with pupil diameter (Supplementary Fig. 2); units that increased firing rate with pupil dilation (40.3% of units, n = 209/519), units that decreased firing rate with pupil dilation (26.0% of units, n = 135/519), and units that showed no significant correlation with pupil diameter (33.7% of units, n = 175/519). Such diversity in PPC spiking activity related to arousal is in general agreement with work in mouse visual and frontal cortices, which found that subpopulations of neurons in mouse visual cortex either increased or decreased firing rate with arousal (Garcia-Junco-Clemente et al., 2017; Vinck et al., 2015). In contrast to PPC, the majority of multi-units in LP/Pulvinar displayed increased firing rate with pupil diameter (72.2%, n = 122/169), with only a negligible portion of units displaying a decrease in firing rate (3.0%, n = 5/169). Together, these results suggest greater heterogeneity of spiking dependence on arousal in cortex, and may reflect the greater complexity and diversity of neuronal populations that comprise cortical circuits (Garcia-Junco-Clemente et al., 2017).

**Figure 2 /.**
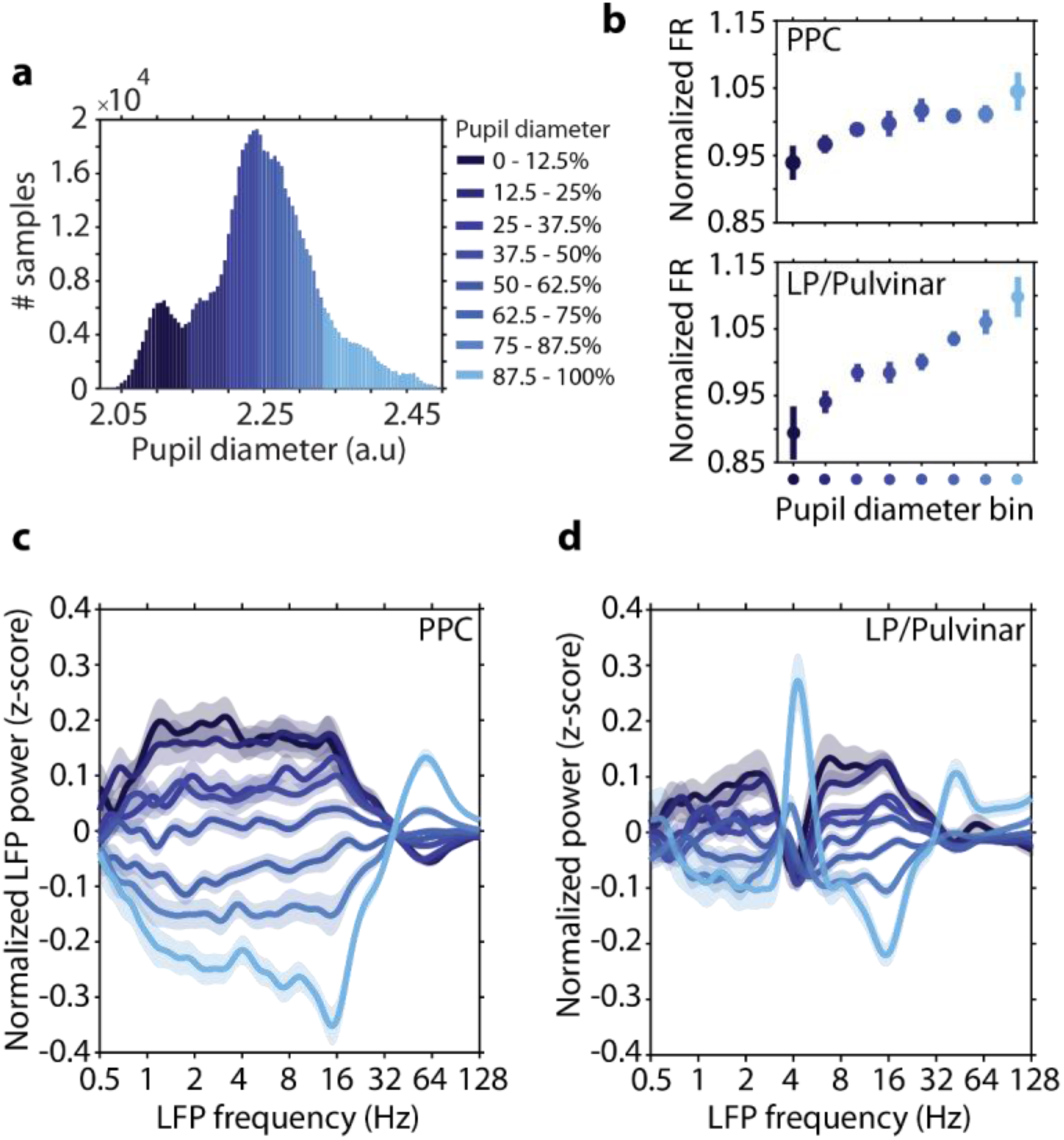
Neuronal spiking rate and LFP spectral power in PPC and LP/Pulvinar are modulated with pupil diameter. **a**, A representative example of how pupil diameter time series were divided up into bins (each bin represents 12.5% of all samples). Neurophysiological data were then analyzed according to pupil diameter bin. **b,** The normalized spiking rate in both PPC and LP/Pulvinar increases with pupil dilation. Color denotes pupil diameter, as indicated in **a**. **c,** The mean (± SEM) z-scored LFP power in PPC as a function of carrier frequency across pupil diameter bins. Low (< 30 Hz) and high (> 30 Hz) frequency LFP power displayed opposing relationships to pupil diameter. **d,** the same as **c** but for LP/Pulvinar LFP power. Note the antagonism between low and high frequency LFP oscillatory power is similar to PPC, with the exception of the theta band (∼4 Hz), which displays increased power during large pupil diameter states.

To examine how signatures of local network dynamics were altered by arousal, we computed changes in LFP power spectra as a function of pupil diameter (Figure 2c, d). Pupil diameter-related changes in LFP power were characterized by opposing effects on low (< 30 Hz) and high frequency (> 30 Hz) LFP oscillations in PPC (for raw power spectra, see Supplementary Fig. 3); low frequency power was stronger during small pupil diameter epochs, while high frequency power was stronger during large pupil diameter epochs (Figure 2c). LP/Pulvinar displayed similar pupil size-dependent antagonism between low and high frequency LFP signals, with the exception of LFP power in theta band (3.3-4.5 Hz), which displayed peak power during large pupil diameter epochs (Figure 2d). In both PPC and LP/Pulvinar, large pupil diameter epochs were associated with reduced LFP power in the 12-17 Hz frequency range. Since LFP signal power and thalamo-cortical coherence in 12-17 Hz frequency range was reduced during visual stimulation (Supplementary Fig. 3), we posit that rhythms in this frequency band represents a homologue of the alpha oscillation in the ferret.

**Figure 3 /.**
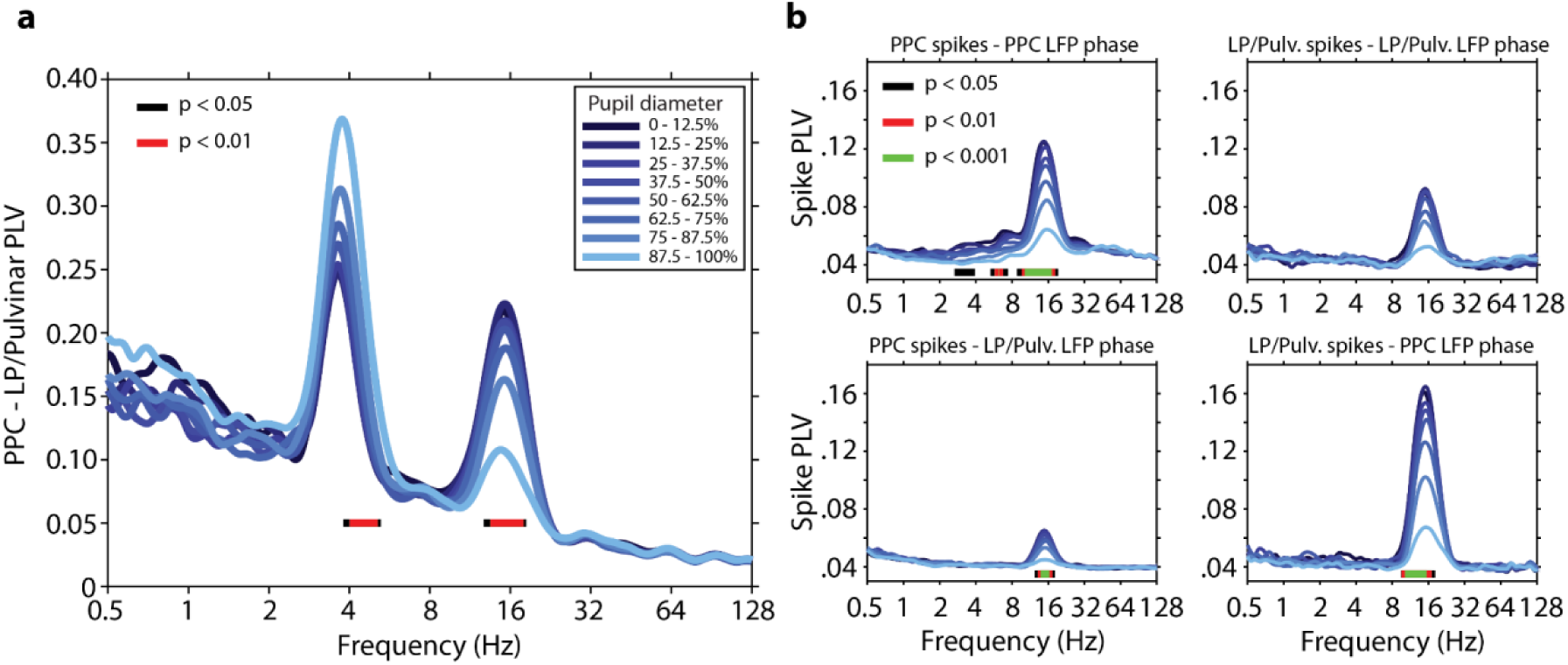
Thalamo-cortical synchronization varies with ongoing fluctuations in pupil-linked arousal. **a,** PLV measured between LP/Pulvinar and PPC as a function of LFP frequency and pupil diameter. Pupil diameter is denoted by color (see legend). Note the prominent thalamo-cortical phase synchronization in the theta (∼4Hz) and alpha (12-17Hz) carrier frequency bands. Significant modulation across pupil diameter bins is indicated by bars plotted below PLV traces (one-way ANOVA, FDR-corrected P-values). PLV in the theta band significantly increased with pupil dilation, while alpha PLV significantly decreased with pupil dilation. **b,** The phase synchronization (PLV) of spiking activity to LFP rhythms recorded within the same brain structure (top row), as well as between regions (bottom-row). Spike-PLV both locally within PPC and LP/Pulvinar, as well as between PPC and LP/Pulvinar revealed phase locking of spiking activity to alpha oscillations. Significant modulation of spike-PLV across pupil diameter bins is indicated by color bars plotted below PLV traces (one-way ANOVA, FDR-corrected P-values). Note that the strongest alpha band spike-PLV was observed between LP/pulvinar spikes and PPC LFP phase.

### Arousal dependent modulation of thalamo-cortical synchronization

Considering that complex behaviors such as attention require the coordination of neural activity between cortex and thalamus (Saalmann et al., 2012), we next asked if signatures of thalamo-cortical functional interaction varied with ongoing fluctuations in pupil-linked arousal. We observed thalamo-cortical LFP phase synchronization in the theta and alpha carrier frequency bands (Figure 3a, Phase delay in theta band = 9.6ms, 0.55 circular variance; Phase delay in alpha band = 30.2ms, 0.78 circular variance). Strikingly, the strength of phase synchronization in these frequency bands was modulated in opposing directions by changes in pupil diameter, with theta phase synchronization significantly increasing (P < 0.01 one-way ANOVA), and alpha phase synchronization significantly decreasing with pupil dilation (Fig. 3a, P < 0.01 one-way ANOVA; see Supplementary Fig. 4 for power-matched PLV analysis).

These results suggest that fluctuations in pupil-linked arousal coincide with a shift in the carrier frequency of thalamo-cortical functional interaction from the alpha band in low arousal states, to the theta band in high arousal states. In line with this hypothesis, spike cross-correlation analysis uncovered synchronous oscillatory patterns of thalamo-cortical neuronal firing occurring in the alpha band (Supplementary Fig. 5). Transition from small to large pupil diameter was marked by a significant decrease in oscillatory spike correlations in the alpha band (Supplementary Fig. 5c, P = 4.1^-6^ one-way ANOVA, r = -0.42). In addition to LFP-LFP and spike-spike correlations, we also observed spike-LFP phase synchronization in the alpha frequency band both locally within LP/Pulvinar and PPC, as well as between thalamus and cortex (Fig. 3b). Spike-LFP phase synchronization in the alpha band significantly decreased with pupil dilation locally within PPC, as well as between regions (P < 0.001 one-way ANOVA). Consistent with previous work in humans(Bonnet and Arand, 2001), peak alpha frequency significantly increased in PPC with arousal (Supplementary Fig. 6, P = 0.003 one-way ANOVA, r = 0.32), however LP/Pulvinar displayed no such relationship. In contrast to results in the alpha band, spike-spike correlation and spike-LFP phase synchronization were weak in the theta frequency band both locally and between regions (Fig. 3b). Together, these results suggest that alpha rhythms synchronize thalamo-cortical network activity in low arousal states, with increasing arousal leading to the desynchronization of spiking activity and a transition to the theta carrier frequency for thalamo-cortical functional interaction.

### Arousal dependent thalamo-cortical effective connectivity

Given that we observed two distinct carrier frequencies of thalamo-cortical functional interaction that were differentially modulated by arousal, we asked if activity in these frequency bands reflected directed interaction between LP/Pulvinar and PPC. We quantified directionality in thalamo-cortical interaction by computing spectrally resolved Granger causality and performing phase slope index (PSI) analyses between LP/Pulvinar and PPC LFP signals (Fig. 4a; see Supplementary Fig. 7 for PSI analysis). In line with previous results (Saalmann et al., 2012), we found significant reciprocal causal interaction between PPC and LP/Pulvinar in the alpha frequency band (P < 0.001, permutation test). However, in contrast to a previous report (Saalmann et al., 2012), we found that the cortical influence on thalamus was significantly stronger than thalamic influence on cortex in the alpha band (P = 0.003, t-test). PSI analyses confirmed this result (Supplementary Fig. 7), with a larger proportion of cortical channels driving thalamus (32%) than vice versa (11%). Both Granger causality and PSI findings are consistent with the hypothesis that alpha rhythms arise from reciprocal interaction between thalamic and cortical alpha oscillators. In contrast to alpha rhythms, Granger causal interaction in the theta band was only observed from LP/Pulvinar to PPC (P < 0.001 permutation test, Fig. 4a). PSI analyses agreed with this result, with 19% of channel pairs displaying significant thalamus-to-cortex interaction, while only 3% displayed significant cortex-to-thalamus interaction (Supplementary Fig. 7). Collectively, these results suggest that theta rhythms predominantly propagate from thalamus to cortex, while alpha rhythms propagate principally from cortex to thalamus.

**Figure 4 /.**
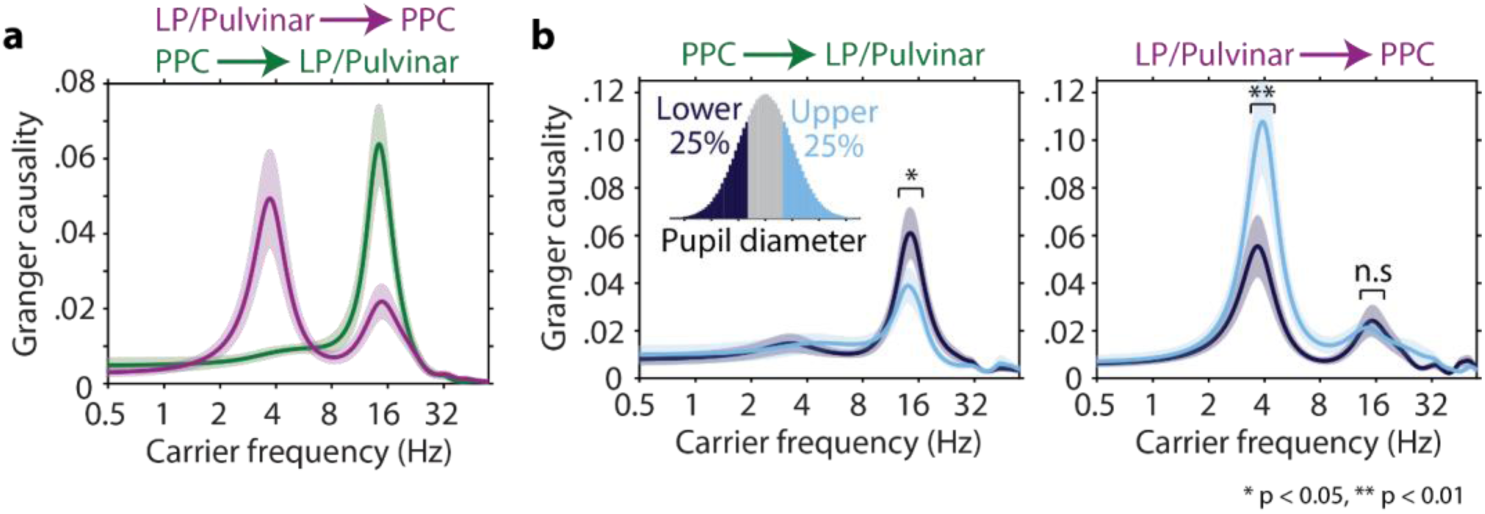
Arousal level determines the direction and carrier frequency of thalamo-cortical causal interaction. **a,** Spectrally resolved Granger causality shows the carrier frequencies of directed interaction from LP/Pulvinar to PPC (magenta), and PPC to LP/Pulvinar (green, ± SEM). LP/Pulvinar has a causal influence on PPC in the theta and alpha frequency bands. In the opposing direction, PPC has a causal influence on LP/Pulvinar in the alpha band. **b,** Granger causality was measured for time periods where the pupil diameter was small (< 25%, dark blue), and large (> 25%, light blue), respectively. Data were subsampled to match power distributions between conditions. The causal influence of PPC alpha oscillations on LP/Pulvinar was significantly stronger during small pupil diameter epochs (left plot, P = 0.018 t-test). In contrast, the causal influence of LP/Pulvinar theta oscillations on PPC was significantly greater during large pupil diameter epochs (right plot, P = 0.0015 t-test).

To examine how thalamo-cortical effective connectivity is modulated by arousal, we recomputed Granger causality for epochs that represented the lower and upper 25% of pupil diameter for each recording, respectively. Since LFP power was highly dependent on arousal, we subsampled data to match power distributions across pupil diameter bins (Supplementary Fig. 8). The causal influence of PPC on LP/Pulvinar in the alpha band was significantly stronger for small pupil diameter states (P = 0.018 t-test, Fig. 4b left). In contrast, LP/Pulvinar causal influence on PPC in the theta band was significantly stronger in large pupil diameter states (P = 0.0015 t-test, Fig. 4b right). These results indicate that fluctuations in pupil linked arousal result in a dynamic switch in the predominant carrier frequency of thalamo-cortical communication and the direction of causal interaction between LP/Pulvinar and PPC.

### Thalamo-cortical network dynamics during visual processing

Until now, we have examined how variability in thalamo-cortical network dynamics relate to ongoing fluctuations in pupil-linked arousal in the absence of visual input. Are such patterns of thalamo-cortical network dynamics exclusive to recordings in the dark, or are they reflective of more generalizable network states that also emerge during periods when animals are engaged in sensory processing? To answer this question, we investigated how patterns of activity in LP/Pulvinar and PPC are modulated in response to the presentation of naturalistic images or videos (Figure 5a). Naturalistic visual stimuli elicited increased gamma power and decreased alpha power in both LP/Pulvinar and PPC (Figure 5b-c for videos; see Supplementary Fig. 9 for images). In general, naturalistic video stimuli elicited more robust spiking and LFP responses than static images (PPC static image firing rate = 13.45 ± 1.65, video firing rate = 14.33 ± 1.64, P = 8.8^-5^, sign test; LP/Pulvinar static image firing rate = 8.72 ± 1.04, video firing rate = 9.68 ± 1.09, P = 8.0^-7^, sign test). In the prestimulus period we found thalamo-cortical LFP synchronization at theta and alpha carrier frequencies (Figure 5d), similar to small pupil diameter states in the dark. However, upon presentation of video stimuli, thalamo-cortical synchronization rapidly shifted to the theta carrier frequency, with an accompanying reduction in alpha synchronization. Theta synchronization maintained throughout the duration of stimulus presentation, before returning to the theta and alpha carrier frequencies after stimulus offset (Figure 5d). Granger causality analysis confirmed that video stimuli induce a rapid reversal in the direction and carrier frequency of thalamo-cortical effective connectivity similar to what we observe during small and large pupil diameter epochs in the dark (Figure 5e). These findings suggest that during presentation of video stimuli the LP/Pulvinar – PPC network shifts towards a state defined by thalamus driven theta oscillations. In addition, these results illustrate that arousal dependent thalamo-cortical network states that spontaneously occur in the dark are also invoked when animals are processing sensory information.

**Figure 5 /.**
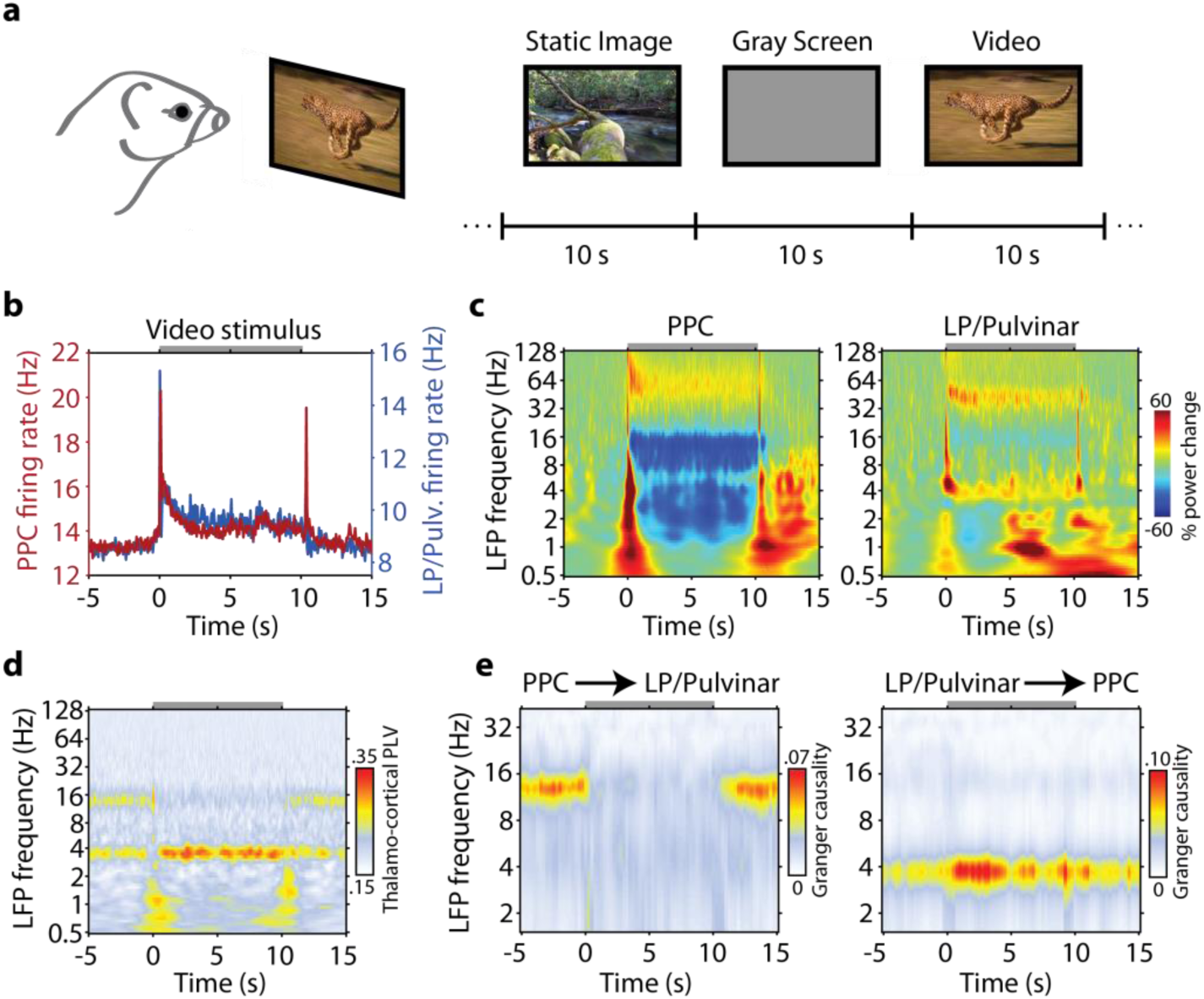
Visual processing induced changes in thalamo-cortical network dynamics. **a,** Animals passively viewed a collection of naturalistic images or videos. During the inter stimulus interval a gray screen was presented. **b**, Population mean spike rate in PPC and LP/Pulvinar during presentation of video stimuli. The gray bar at the top of the plot indicates the duration of stimulation. **c**, Population LFP spectrograms from PPC (left) and LP/Pulvinar (right) during presentation of naturalistic videos. LFP power was normalized to the period -5 to -1 seconds before stimulus onset. Note the decrease in alpha oscillatory power in PPC during stimulus presentation. **d**, Across session average thalamo-cortical phase synchronization in response to naturalistic video stimuli. PLV in the theta band is elevated during stimulus presentation, while alpha PLV is weaker. **e**, Time and frequency resolved Granger causality analysis computed between PPC and LP/Pulvinar LFP signals for naturalistic video stimuli. The onset of video stimuli leads to a breakdown of PPC causal influence on LP/Pulvinar in the alpha band (left plot), and an increase of LP/Pulvinar causal influence on PPC in the theta frequency band (right plot).

### Thalamo-cortical network dynamics and pupil-linked arousal correlate with saccade behavior

What is the link between the emergence of these patterns of thalamo-cortical network dynamics and behavioral output? Given the well-established link between saccadic eye movements and cognitive processes (Liversedge and Findlay, 2000), we answer this question by examining how thalamo-cortical network dynamics relate to saccade behavior. Saccade kinetics in the ferret displayed similar characteristics to saccades in humans (Otero-Millan et al., 2008) and non-human primates (Bosman et al., 2009) (Figure 6a, Supplementary Fig 10), albeit with a lower overall saccade rate (0.32 ± 0.03 Hz). Animals displayed an elevated rate of saccades while viewing video stimuli (0.35 ± 0.03 saccades/s videos stimuli, 0.19 ± 0.02 saccades/s prestimulus, P = 2.7^-7^ t-test, Figure 6b), illustrating that ferrets utilize saccades to actively sample the visual environment.

**Figure 6 /.**
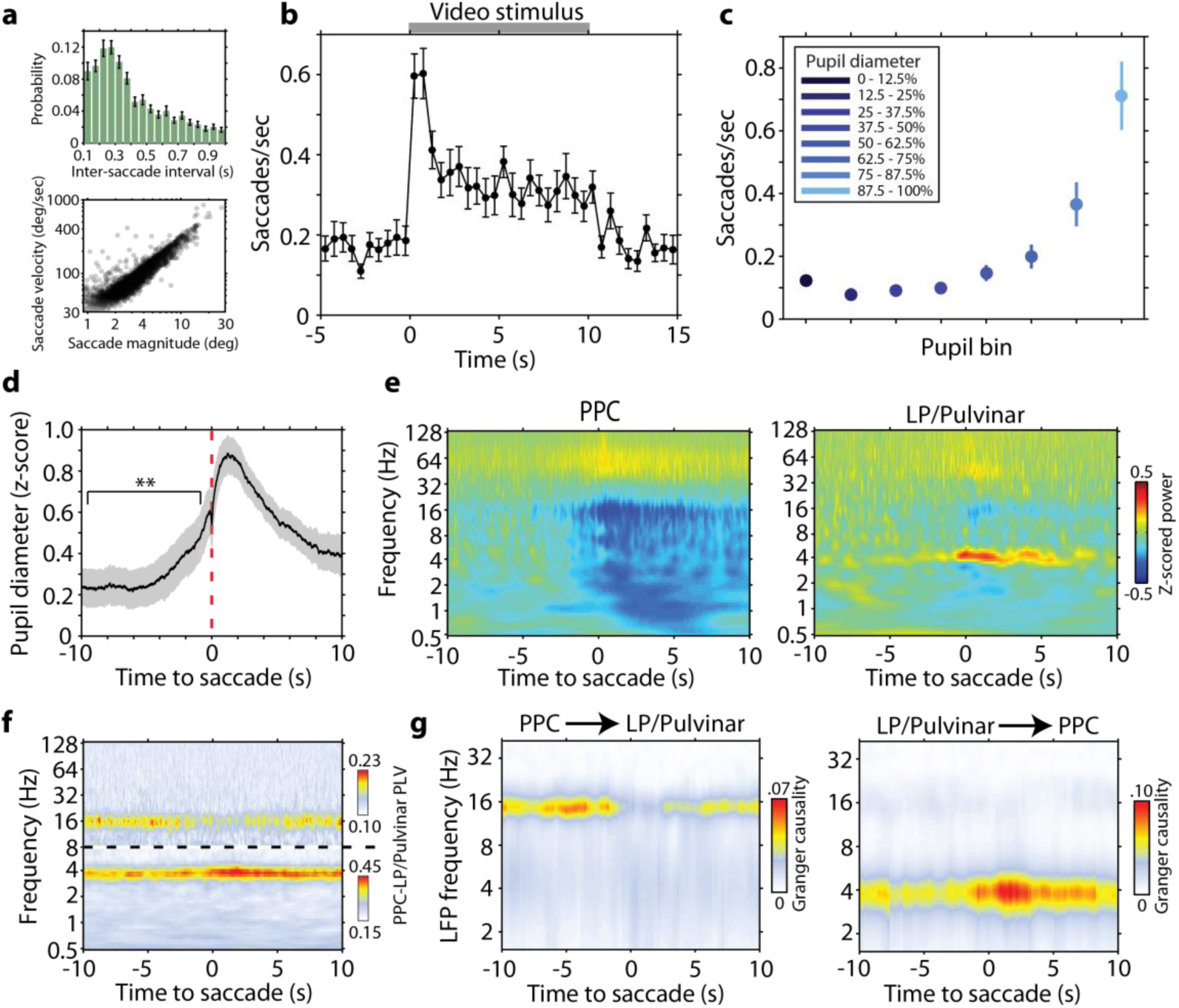
Saccades link visual sensory processing and pupil-linked arousal related changes in thalamo-cortical dynamics. **a,** Distribution of inter saccade interval (top, ± SEM) shows that ferrets actively sample the visual environment rhythmically. The relationship between saccade magnitude and peak velocity (bottom) reflects the ballistic nature of saccades in ferrets. **b**, Saccade rate during naturalistic video presentation illustrates that animals actively sample visual stimuli by showing an elevated rate of saccades (n = 26 sessions). **c**, Saccade rate as a function of pupil diameter in the dark. The rate of saccades during large pupil diameter states is comparable to the rate of saccades when animals are actively sampling naturalistic videos. **d**, Mean (± SEM) fluctuations in pupil diameter time locked to saccadic eye movements in the dark. Transient increases in pupil diameter precede saccades (P = 0.004, t-test). **e**, Population average LFP power spectrograms in PPC (left) and LP/Pulvinar (right) time locked to saccades in the dark. LFP power was z-score normalized across the entire recording session. **f**, Population average PPC to LP/Pulvinar PLV time locked to saccades in the dark. Dotted line indicates a break in the color scale, as shown to the right of the figure. **g**, Across session average time and frequency resolved Granger causality between PPC and LP/Pulvinar around the occurrence of saccades in the dark. Active sampling by saccades was associated with a decrease in PPC causal influence on LP/Pulvinar in the alpha band and an increase in LP/Pulvinar on PPC in the theta band.

Animals also performed saccades in the dark (Supplementary Fig. 10), where saccade rate was significantly modulated by ongoing fluctuations in pupil diameter (Figure 6c, P = 3.80^-15^ one-way ANOVA). Indeed, saccade-triggered analysis of pupil diameter in the dark revealed that the pupil significantly dilated prior to the onset of saccadic eye movements (P = 0.004 t-test, Figure 6d). This result suggests that oculomotor behavior may arise from a more general shift towards an aroused state. In line with this, PPC and LP/Pulvinar LFP power during saccades closely resembled power spectra for large pupil diameter states (Figure 2c-d), with power modulations occurring over a time course of several seconds around saccades (Figure 6e). Similarly, we observed a decrease in thalamo-cortical synchronization in the alpha band coinciding with saccades accompanied by a matching increase in theta band synchronization occurring over a slower timescale (Figure 6f).

To test if the direction of thalamo-cortical interaction also changed during saccade behavior, we computed spectrally resolved Granger causality analysis time-locked to saccades (Figure 6g). Saccades were associated with a significant decrease in PPC causal influence on LP/Pulvinar in the alpha band, accompanied by a significant increase of LP/Pulvinar causal influence on PPC in the theta band (P < 0.05, test against random saccade times, Supplementary Figure 11). These results strongly support the hypothesis that dynamic changes in thalamo-cortical causal interaction facilitate active visual sampling of the environment.

Recent work has shown that faster pupil dilations are correlated with noradrenergic input to cortex, whereas slower fluctuations are correlated with cholinergic input(Reimer et al., 2016). Consistent with a role for noradrenergic input in modifying thalamo-cortical network dynamics, we found striking shifts in spike rate, LFP power and saccade rate time locked to the occurrence of transient pupil dilations (Supplementary Fig. 12).

How do ongoing fluctuations in thalamo-cortical network dynamics affect the way animals sample visual stimuli? To answer this question, we correlated the number saccades performed during each trial of visual stimulus presentation with the strength of thalamo-cortical synchronization in the theta and alpha frequencies bands (Figure 7a). We found a significant positive correlation between synchronization in the theta band and the number of saccades performed per trial for both video and image stimulus conditions (Figure 7a for video stimuli, P = 0.0002 R = 0.17; for image stimuli see Supplementary Fig. 13, P = 0.0009 R = 0.15). In contrast, we found a significant negative correlation between synchronization in the alpha band and the number of saccades per trial (video stimuli P = 0.011 R = -0.11; image stimuli P = 0.0001 R = -0.17). Thus, the state of the thalamo-cortical system as defined by its oscillatory functional connectivity indexed the level of engagement with the external world.

**Figure 7 /.**
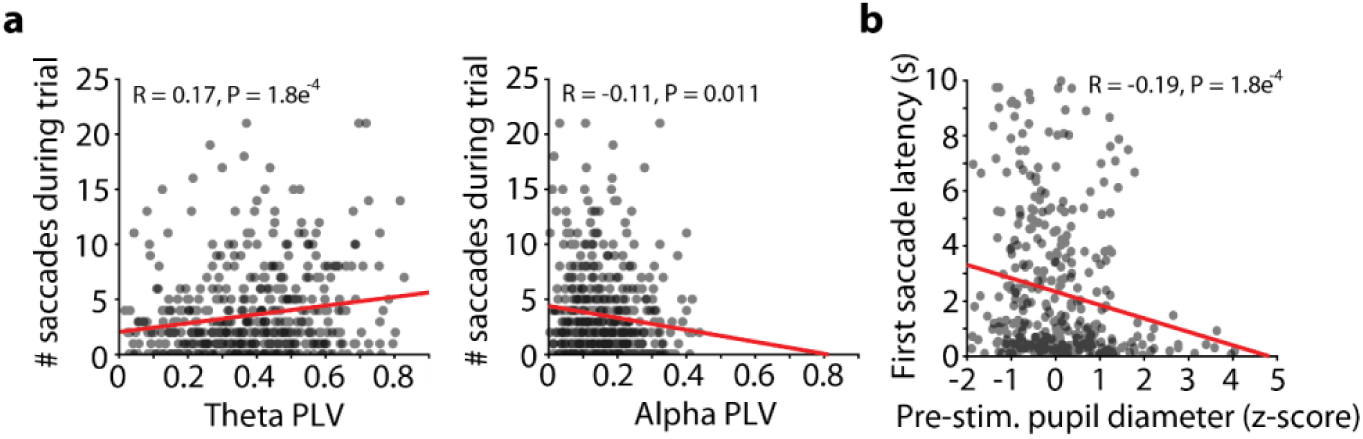
Thalamo-cortical synchronization and pupil-linked arousal correlate with saccade behavior. **a,** Correlation of thalamo-cortical phase synchronization in the theta (left) and alpha (right) carrier frequency bands and the number of saccades performed during presentation of naturalistic video stimuli. Theta PLV displays a significant positive correlation with saccadic sampling of stimuli, whereas alpha PLV displays a significant negative correlation. **b**, Correlation of pre-stimulus pupil diameter to the latency of the first saccade for subsequent naturalistic video stimulus presentation. Significant negative correlation illustrates that animals react by visually sampling stimuli more rapidly when they are in a more aroused state.

Finally, to show that fluctuations in pupil-linked arousal also affect how animals process incoming sensory information, we correlated the prestimulus pupil diameter with the latency of the first saccade for each trial. We found a significant negative correlation between prestimulus pupil diameter and first saccade latency for both image and video stimuli (Figure 7b, P = 0.0001 R = -0.19 for video stimuli; for images see Supplementary Fig. 13b, P = 6.39^-5^ R = -0.22), indicating that animals sampled more rapidly when incoming sensory input arrived during a state of heightened arousal. Taken together, these results show that ongoing fluctuations in pupil-linked arousal and associated changes in thalamo-cortical functional interaction affect the way in which animals sample the external environment with saccadic eye movements.

### Discussion

Due to their reciprocal connectivity with widely distributed cortical areas, higher-order thalamic structures are proposed to play an important role in orchestrating patterns of large-scale cortico-cortical and thalamo-cortical interaction that underlie cognition(Saalmann and Kastner, 2011). Our results provide the first evidence that pupil-linked arousal, and by extension neuromodulation, play an important role in dynamically sculpting these patterns of thalamo-cortical functional interaction. We demonstrate that ongoing fluctuations in pupil-linked arousal lead to dynamic switching of both the direction and carrier frequency of thalamo-cortical communication, with low arousal states marked by cortical alpha oscillations driving synchronized activity between LP/Pulvinar and PPC, while higher arousal states were marked by LP/Pulvinar driving PPC in the theta frequency band. Furthermore, we show similar transitions in thalamo-cortical network dynamics during visual processing, and in particular, during active sampling of the external environment via saccades.

How does cortically driven alpha synchronization in thalamo-cortical networks facilitate the computations that underlie cognition in states of low arousal? One prominent hypothesis is that alpha oscillations reflect the precise temporal parsing of cortical activity via phasic inhibition (Klimesch, 2012). Under this framework, layer 5 projection neurons in PPC entrain local alpha oscillations in the LP/Pulvinar through pulsed volleys of action potentials. Given that LP/Pulvinar projections to superficial layers of early visual cortex influence cortical state (Purushothaman et al., 2012; Roth et al., 2016), PPC to LP/Pulvinar synchronization in the alpha band may act to gate or suppress the processing of incoming sensory information at early stages of visual cortex. Beyond this gating role, alpha oscillations have also been associated with cognitive processes that require internalization of attention, such as working memory (Klimesch, 2012; Roux and Uhlhaas, 2014) and creativity (Fink and Benedek, 2014). In agreement with the internalization of attention, we consistently observed reduced saccade behavior during periods of elevated alpha oscillations. Similar to humans (Adrian, 1934), this alpha-dominant mode of network dynamics was disrupted by the presentation of visual stimuli. Interestingly, we found that the degree to which animals actively sampled stimuli with saccades was negatively correlated with thalamo-cortical synchronization in the alpha band. Thus, the dynamic switching away from the synchronized alpha oscillation network state may be a signature of the transition between internalized and overt processes of attention.

We note that the endogenous alpha frequency identified here does not match the classic 10 Hz rhythm reported in humans. Indeed, we favor a definition of neural rhythms based on underlying physiological mechanisms, as opposed to the arbitrary assignment of bands based on carrier frequency. Given that alpha oscillations are proposed to arise through reciprocal thalamo-cortical interaction (Bollimunta et al., 2011), it is unsurprising that these rhythms exhibit a shorter period in animals with smaller brains (and therefore shorter conduction delays), such as dogs (da Silva et al., 1973), cats (Hughes et al., 2004), and ferrets (Stitt et al., 2015).

In contrast to states of low arousal, thalamus-driven theta oscillations during states of high arousal were associated with an increase in saccade behavior. We found that transitions between alpha- and theta-dominant modes of thalamo-cortical synchronization arose transiently around the occurrence of saccades. These data suggest that the causal influence of thalamus on cortex in the theta band may play a role in active visual sampling and overt attention. Lending support to this, previous studies in primates found that cortical LFP signals synchronize to microsaccades occurring rhythmically at the theta frequency (Bosman et al., 2009). Furthermore, theta oscillations temporally coordinate gamma band synchronization between lower and higher order visual cortices during attention allocation (Bosman et al., 2012). Therefore, the thalamic causal influence on cortex in the theta band may represent a mechanism to selectively synchronize distributed visual cortical regions (Jones, 2001; Saalmann et al., 2012), facilitating visual exploration and attentional selection. Given that LP/Pulvinar is reciprocally connected with widely distributed visual cortical regions, it certainly exhibits the anatomical framework to play such a role in orchestrating cortical network dynamics and information routing (Jones, 2001). However, we observed only weak locking of spiking activity to theta oscillation phase in LP/Pulvinar, suggesting the inputs that synchronize LP/Pulvinar and PPC theta oscillations are likely subthreshold and do not predominantly originate from the population of LP/Pulvinar neurons we recorded from. Alternatively, these theta-synchronizing inputs may originate from other brain regions such as prefrontal cortex (Saalmann, 2014).

Fluctuations in pupil diameter under constant luminance have been used as a general measure of arousal and cognitive load (Laeng et al., 2012). A common assumption regarding the relationship between pupil diameter and arousal was that the locus coeruleus (LC), the main source of noradrenergic neuromodulatory input to the forebrain, must somehow form part of the brainstem circuit that controls pupil motility. Indeed, several invasive electrophysiological studies in monkeys have shown that fluctuations in pupil diameter under constant luminance are correlated to neural activity in the LC (Aston-Jones and Cohen, 2005; Joshi et al., 2016). However, correlations were generally weak, and no direct anatomical connection linking the LC and brainstem pupil motility nuclei could be found to explain such correlations (Nieuwenhuis et al., 2011; Wang and Munoz, 2015). In a recent study Reimer et al (2016) showed that fluctuations in pupil diameter track both noradrenergic and cholinergic input to cortex, with more rapid pupil dilations reflecting changes in noradrenergic input, and longer-lasting fluctuations reflecting the sustained activity of cholinergic synapses. Considering that we observe strong transitions in activity patterns and saccade behavior time locked to transient pupil dilation events (Supplementary Fig. 12), we postulate that noradrenergic neuromodulation plays a crucial role in shaping thalamo-cortical network dynamics and oculomotor behavior.

How do ongoing fluctuations in neuromodulatory tone alter large-scale network interactions in the brain? Previous in vitro work has classified in detail the effects of various neuromodulatory subsystems on intrinsic cellular properties (McCormick, 1989; McCormick et al., 1993; Pape and McCormick, 1989; Rogawski and Aghajanian, 1980; Salgado et al., 2016) and the generation of rhythmic activity in local circuits (Lorincz et al., 2008). However, these local cellular changes do not explain the dynamic rerouting of thalamo-cortical communication that we observe at the network level. Experimental evidence relating pupil-linked arousal or neuromodulatory tone to the dynamic reorganization of network-level interaction is sparse. Recent data has shown that pharmacological blocking of noradrenaline reuptake in humans leads to a network-specific reduction in the correlation of hemodynamic signals between brain regions (van den Brink et al., 2016). These findings are consistent with our results on the pupil-linked arousal related reduction in thalamo-cortical spike correlations.

Perhaps the most compelling mechanistic description of how neuromodulation shapes network interactions has come from computational modeling and theoretic studies of complex networks. Most recently, Heeger (2017) proposed a computational framework of cortical processing where network dynamics emerge from the interaction of feedforward and feedback inputs, as well as prior (expectation) driving factors. In this model, a number of state parameters modify the relative contribution of feedforward and feedback processing stages toward the predominating network dynamic. Our findings support Heeger’s hypothesis that such state parameters represent the various neuromodulatory subsystems. Furthermore, Kirst et al (2016) showed that alteration of low-level features of complex networks not only leads to perturbations in the collective dynamics of the network, but can also result in the self-organized rerouting of information flow between network modules. Therefore, although neuromodulators act primarily on intrinsic cellular properties of neurons in cortex and thalamus, when translated to the network level, these local changes may lead to emergent shifts in thalamo-cortical functional interaction.

Although we studied the LP/Pulvinar – PPC thalamo-cortical network here, we speculate that our findings on arousal-dependent changes in interaction may generalize to other thalamo-cortical networks, and perhaps even modes of cortico-cortical functional interaction. One intriguing possibility is that subpopulations of neurons in the LC or basal forebrain differentially modulate the dynamics of specific sub-networks in the brain. Indeed, the heterogeneous nature of LC neuronal activity (Totah et al., 2017) coupled with non-overlapping projection patterns of ascending neuromodulatory fibers (Chandler et al., 2014) may enable such network specific modulation of functional interaction.

Extending these results further, we speculate that deficiencies in the neuromodulatory control of large-scale network interaction may represent a key mechanism underlying the pathology of various neuropsychiatric disorders (Uhlhaas and Singer, 2012). A broader understanding of how neuromodulators shape functional interaction within affected brain networks will enable the targeted design of therapies aimed at restoring physiological patterns of network dynamics.

## Acknowledgements

We thank V. Crunelli, T. Pfeffer, A. Urai, K. Sellers, and Y. Li for comments on the manuscript; C. Yu for technical advice on obtaining recordings from LP/Pulvinar in ferrets; and G. Nolte for advice on data analysis.

### Author Contributions

I.S. and F.F. conceived and designed experiments. I.S. and Z.C.Z. performed implantation of microelectrode arrays, recorded data, and performed anatomical tracing experiments. S.R.S interpreted anatomical data. I.S analyzed pupil and electrophysiological data. I.S., Z.C.Z., and F.F. wrote the paper. All authors contributed to the discussion of the results and revision of the manuscript.

## Methods

### Animals

Five adult spayed female ferrets (*Mustela putorius furo*) were used in this study. Animals had ad libitum access to food pellets and water, and were group housed in cages under standard ambient conditions (12 hour day/light cycle). All animal procedures were performed in compliance with the National Institutes of Health guide for the care and use of laboratory animals (NIH Publications No. 8023, revised 1978) and the United States Department of Agriculture, and were approved by the Institutional Animal Care and Use Committee of the University of North Carolina at Chapel Hill.

### Headpost and electrode implantation

Animals were initially anesthetized with an intramuscular injection of ketamine/xylazine (30mg/kg of ketamine, 1–2mg/kg of xylazine). After loss of the paw pinch reflex, animals were intubated to enable artificial ventilation and delivery of isoflurane anesthesia (0.5-2% isoflurane in 100% oxygen). Throughout surgical procedures, physiological parameters such as the electrocardiogram, end-tidal CO_2_, partial oxygen concentration, and rectal temperature were constantly monitored to maintain the state of the animal. All surgical procedures were performed under sterile conditions. To enable the accurate planning of LP/Pulvinar electrode penetrations, animals were fixed into a stereotactic frame with stainless steel ear bars. The skull was then tilted such that it was oriented perpendicular to the surface of the surgical table. The skin and muscle were reflected to expose the surface of the skull. A custom-designed stainless steel headpost was firmly secured to the anterior extent of the exposed skull with bone screws. A craniotomy was performed on the left hemisphere to expose PPC(Manger et al., 2002) and the cortex overlying LP/Pulvinar. The dura was carefully removed, before lowering a multielectrode array into LP/Pulvinar along stereotactic coordinates(Yu et al., 2016) (2x8 tungsten electrodes, 250μm spacing, 9mm length, Microprobes for Life Science). A second multielectrode array (4x8 tungsten electrodes, 200μm spacing, Innovative Neurophysiology) was then inserted into the deep cortical layers of PPC. Both microelectrode arrays had reference electrodes that were directly adjacent to recording electrodes in the brain. In 3 animals, the PPC electrode was placed into the lateral gyrus, while in 1 animal the PPC electrode was placed in the suprasylvian gyrus. Multielectrode arrays were fixed in place with dental cement. Skin and muscle around the incision was then sutured together. After surgery, animals were administered preventative analgesics and antibiotics for one week. Animals were allowed to recover in their home cage for at least one week prior to recordings.

### Anatomical tracing experiments

One adult female ferret was used for anatomical tracing. Preparation of the animal for aseptic surgery was performed according to procedures described above. Craniotomies were drilled above the PPC on both hemispheres and the dura was removed to expose the underlying cortical surface. A total of 0.8μL of anterograde tracer (rAAV5-CaMKII-mCherry; UNC Vector Core) was injected between depths of 800 μm and 400 μm below the cortical surface in the left hemisphere. The location of the injection matched coordinates of cortical microelectrode array implantation in animals used for electrophysiology experiments (Supplementary Fig. 1). Retrograde tracer (0.8μL cholera toxin subunit B conjugated to Alexa 488; Thermo Fisher Scientific) was injected at the corresponding location in the right hemisphere. Kwik-kast (World Precision Instruments) was applied to the cortex to seal the craniotomy, before a layer of dental cement was applied to prevent regrowth of tissue over the injection sites. Three weeks following tracer injection surgery, the animal was humanely euthanized with an overdose of ketamine/xylazine, and then perfused with 0.1 M PBS initially, followed by 4% paraformaldehyde solution in 0.1 M PBS. After two days of fixation in 4 % paraformaldehyde, the brain was transferred to 30% sucrose in 0.1 M PBS solution for cryoprotection. The brain was then sectioned into 50 μm slices using a cryostat (CM3050S, Leica Microsystems). Sections were imaged on a Nikon Eclipse 80i widefield microscope, with green, red, and brightfield images overlaid to construct composite images to illustrate fluorescent labeling in PPC and LP/Pulvinar.

### Recording procedure

After recovery from surgery, animals were placed into a custom designed behavioral tube and were head-fixed with a stainless steel headpost clamp. PPC and LP/Pulvinar multichannel recording electrodes were then connected to a data acquisition system (INTAN technologies). An infra-red eye tracking camera was focused on the animal’s right eye (ISCAN ETL-200), with instantaneous measurements of pupil diameter, and pupil center x/y position delivered as voltage signals to the data acquisition system. The infrared light source was positioned in the lower right of the animal’s visual field and emitted non-visible light at a wavelength of 940 nm. Broadband extracellular potentials from both multielectrode arrays and eye tracking data were sampled at 30 kHz. Unless otherwise stated, recordings took place in a completely dark room under constant luminance (0.6 Fc). There was a dark adaptation period of at least 30 seconds prior to the start of each recording. To prevent animals from falling asleep while head-fixed, two experimenters engaged in verbal discourse in front of the animal. Recordings in the dark typically lasted between 8 to 13 minutes. In addition to recordings in the dark, data were collected also while animals passively viewed a library of 20 naturalistic images and 20 videos on a screen placed 28.5cm in front of the animal. The eye tracking camera and infra-red light source did not occlude the animal’s view of any part of the screen. Images and videos were randomly interspersed with a stimulus duration of 10 seconds. A gray screen was presented during the inter stimulus interval (inter-stimulus interval = 10 seconds, Figure 5A). Visual stimuli were presented in a well-lit room (ambient luminance 32.6 Fc). Data were collected from multiple sessions across several days or weeks from each animal (number of sessions per animal: 6, 4, 7, 8).

### Verifying electrode positions with histology

Animals were deeply anesthetized according to methods described above. Electrolytic lesions were then produced on both PPC and LP/Pulvinar probes by passing 5 μA of current (10 second pulse) between selected recording electrodes and the reference electrode on each probe. Animals were perfused and brains sectioned according to procedures described above. Brain slices were washed with 0.1 M PBS and stained for cytochrome oxidase(Wong-Riley, 1979), and then imaged with a widefield microscope (Nikon Eclipse 80i; Nikon Instruments). Electrode positions in thalamus were confirmed by either the location of electrode tracks or electrolytic lesions. Electrodes that were deemed to fall outside of LP/Pulvinar were omitted from analysis (number of electrodes omitted per animal: 0, 2, 8, 6).

### Data analysis

All offline data analyses were performed using custom software in Matlab (Mathworks).

#### Data preprocessing

To extract multi-unit spiking activity from broadband extracellular potentials, we band-pass filtered data between 300 and 5000Hz and applied a threshold at -4 standard deviations. Spikes that were detected on more than 3 channels simultaneously were omitted from further analysis. LFPs were obtained by low pass filtering broadband extracellular potentials at 300 Hz in both the forward and reverse direction to avoid phase shifts (4^th^ order Butterworth filter). LFPs were then downsampled to a sample rate of 1 kHz. Eye position and pupil diameter signals were processed in the same way as the LFP.

#### Pupil diameter

To enable a thorough analysis of neuronal dynamics related to changes in pupil dilation, pupil diameter time series were discretized into eight bins of equal size, such that each bin represented 12.5% of samples from the entire recording (Fig. 2a). If the total number of samples per pupil bin did not exceed 30 seconds of cumulative data, then the recording was discarded (n = 6).

#### Saccade detection

Eye azimuth and elevation signals were low pass filtered at 50 Hz to remove potential noise. Blinks were detected automatically, and data within ± 1 second from blinks were removed from analysis. Elevation and azimuth time series were converted into eye velocity vectors. A threshold was then set at 4 standard deviations of the eye velocity vector to detect saccades. A subset of recording sessions in the dark were excluded from saccade based analyses due to high frequency noise in eye-position signals (n = 12).

#### Spectral decomposition

Time frequency estimates were computed by convolving LFP time series with Morlet wavelets that were Gaussian shaped in both the time and frequency domain(R. Kronland-Martinet, 1987; Tallon-Baudry et al., 1997).

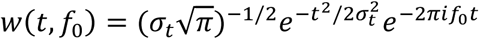

Where *w*(*t, f*_0_) is the complex Morlet wavelet at carrier frequency *f*_0_, and *σ*_*t*_ is the standard deviation of the wavelet in the time domain. *σ*_*t*_ is defined as

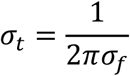

Where *σ*_*f*_ is the standard deviation in the frequency domain, and is defined as a constant depending on the wavelet carrier frequency *σ*_*f*_ = *f*_0_/7. Time frequency estimates *X*(*t, f*_0_) were then computed by convolving LFP time series *x*(*t*) with complex Morlet wavelets *w*(*t, f*_0_).

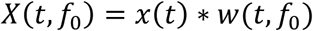

Where * denotes the convolution operation. We convolved LFP signals with a family of 80 Morlet wavelets that had carrier frequencies logarithmically spaced between 0.5 and 128 Hz. The power of LFP signals across all carrier frequencies was computed by taking the absolute value of squared time frequency estimates.

#### LFP phase synchronization

To quantify phase synchronization between simultaneously recorded LFP signals, we computed the phase locking value(Lachaux et al., 1999) (PLV). Briefly, the phase angle between real and complex components of time frequency estimates was calculated for thalamic *θ*^*t*^ and cortical *θ*^*c*^ signals. PLV at the carrier frequency *f*_0_ for *N* samples was then defined by the following formula

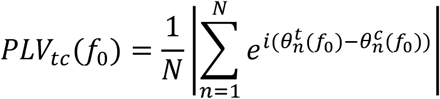

Assuming a uniform circular distribution of phases, the magnitude of *PLV*_*tc*_ is biased towards 1 with few observations, and towards 0 with many observations. To control for this bias, pupil diameter based PLV analyses were computed using a constant number of randomly drawn phase angles between conditions and animals (30000 samples equating to 30 seconds of data). This was repeated for each thalamo-cortical channel pair 100 times, with the mean *PLV*_*tc*_ across all permutations used for further analysis. For each recording session, the average thalamo-cortical *PLV*_*tc*_ between all PPC and LP/Pulvinar channel pairs was calculated, and then used to compute the across-session average PLV.

One potential confound that arises when comparing PLV analyses between conditions with varying spectral power (for example alpha/theta power across pupil diameter bins), is that significant differences in PLV may emerge spuriously due to changes in the signal to noise ratio. To control for any potential signal to noise confounds for PLV estimates, we performed additional analyses where data were subsampled to match the power distributions across pupil diameter bins for each frequency band (Supplementary Fig. 4).

#### Spike-LFP phase synchronization

To determine the dependence of spike timing on LFP oscillation phase, we computed the PLV(Lachaux et al., 1999) between co-recorded spikes and LFPs both within and between brain regions. For within region analysis, spike PLV was computed between all channel combinations on each microelectrode array. Spike PLV analysis was not computed using the LFP on the same electrode to avoid spectral bleeding of spike waveforms contaminating phase synchronization estimates. For between region analyses, spike PLV was computed for all thalamo-cortical channel pairs. The spike PLV at a carrier frequency of *f*_0_ for *K* spikes was defined as

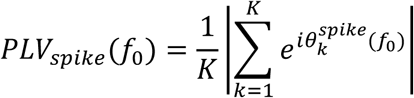

Where *θ*^*spike*^ represents the instantaneous LFP phase at the occurrence of each spike. Spike PLVs were computed from 200 randomly drawn spikes for each channel pair combination. This process was repeated 40 times to obtain an estimate of the mean spike PLV. Channels where less than 200 spikes were detected for each pupil diameter bin were omitted from spike PLV analysis.

#### Granger causality

We computed Granger causality to measure the directed influence thalamic LFPs have on cortical LFPs, and vice versa. Granger causality is rooted in the autoregressive (AR) modeling of time series, where future values of a process *x*(*t*) are modeled based on previous values of *x*(*t*). In this framework, a separate process *y*(*t*) can be said to have a causal influence on *x*(*t*) if the past values of *y*(*t*), when accounted for in a bivariate AR-model, improve the prediction of future values of *x*(*t*) beyond that obtained by the univariate AR-model of *x*(*t*) alone(Granger, 1969). While classically employed in the time domain, the computation of Granger causality can be operationalized in the frequency domain to uncover the physiological carrier frequencies of directed interaction between brain regions(Geweke, 1982). We computed Granger causality between co-recorded LP/Pulvinar and PPC LFP signals in the frequency domain using the multivariate Granger causality (MVGC) toolbox for Matlab(Barnett and Seth, 2014). LFP signals from both regions were low-pass filtered at 100 Hz with a phase preserving filter, and then downsampled to 200 Hz. To reduce dimensionality, we computed representative signals from thalamus and cortex by calculating the median downsampled LFP signal across all channels in PPC and LP/Pulvinar microelectrode arrays. These signals were then windowed into segments of one second (200 samples) length. Model order was then selected based on the minimum Akaike information criterion value(Barnett and Seth, 2014), with a maximum model order of 20 allowed. Vector AR-models were then checked to ensure that they reliably captured the spectral content of input data. Spectral Granger causality was then computed according to routines from the MVGC toolbox. A random permutation test was used determine the significance of Granger causality peaks for individual recording sessions. Data segments from one brain region were randomly shuffled in a procedure that maintained spectral content, while disturbing the temporal codependence of co-recorded LFP signals. Granger causality measured on shuffled data represented the directed interaction that arises by chance based on the spectral signatures of the underlying neural processes. This procedure was repeated 100 times to generate a distribution of thalamo-cortical causal influence expected by chance. The distance of original Granger causality estimates from the shuffled mean were expressed in terms of standard deviation of the shuffled distribution, with values larger than 3 indicating significant Granger causal influence (p < 0.01). For Granger causality analysis based on pupil diameter, recordings were split into segments representing the lowest 25% or highest 25% of the pupil diameter distribution (Fig. 3b). Only data segments where pupil diameter was maintained in low or high states for more than one second were included. To control for signal to noise confounds, we subsampled data to match power distributions between low and high pupil diameter conditions (Supplementary Fig. 8). Power subsampling was performed based on the power of the predominant carrier frequency of thalamo-cortical causal influence in each direction (theta for LP/Pulvinar to PPC, alpha for PPC to LP/Pulvinar). To detect significant differences in the strength of Granger causality between low and high pupil diameter conditions, we performed t-tests on identified spectral peaks from data pooled across all recordings. In addition to pupil diameter linked granger causality, we computed time-resolved Granger causality around the occurrence of saccades with a sliding window of 1 s length and a step size of 0.25 s. To determine the significance of saccade related changes in Granger causality, we recomputed Granger causality at random time points throughout each recording session, where the number of data segments is equal to the number of detected saccades. This was repeated 1000 times for each recording session. Saccade triggered Granger causality spectra were then normalized by the mean and standard deviation of randomly computed Granger causality spectra. Significant time and frequency points were those that deviated from the mean of randomly computed Granger causality estimates by 2 standard deviations (p < 0.05).

#### Phase slope index

Phase slope index (PSI) analysis is a non-autoregressive model based method that was used to quantify effective connectivity between co-recorded thalamic and cortical LFP signals. This form of analysis is grounded in the idea that if one brain region drives another brain region with a constant time lag, then one might expect the relative phase lag between signals in each region to increase as a function of carrier frequency(Nolte et al., 2008). PSI analysis was computed as described previously(Nolte et al., 2008). Briefly, LFP time series were windowed into segments of 1024 samples and the Fast Fourier Transform (FFT) on each data segment was computed. The complex coherency between thalamic and cortical signals at a carrier frequency of *f*_0_ was then defined as

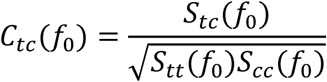

Where *S*_*tc*_(*f*_0_) represents the cross spectra, or the Fourier transform of thalamic signals 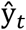 multiplied with the complex conjugate of the Fourier transform of cortical signals 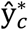

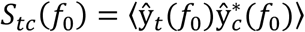

〈 〉 indicates the computation of the expectation value. The phase slope is then computed across the frequency band of interest *f* → *F* as follows

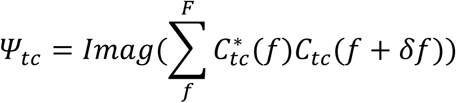

Where ^*^ denotes the complex conjugate and δ*f* denotes the (*f* + 1)^*th*^ resolved frequency of the Fourier transform. The standard deviation of *ψ*_*tc*_ was estimated using the jackknife resampling method. Finally, *ψ*_*tc*_ was normalized by the standard deviation to estimate significance of effective thalamo-cortical connectivity, with values above 2 indicating that thalamus significantly drives cortex, and values below -2 indicating that cortex significantly drives thalamus (p < 0.05). We initially computed PSI using a sliding bandwidth of 4 Hz in the frequency domain to determine the carrier frequencies of thalamo-cortical effective connectivity. Since we observed both significant drivers and receivers across multiple frequency bands, the standard deviation of normalized PSI values was used to quantify the spread of reciprocal thalamo-cortical effective connectivity in the frequency domain. After identifying that the theta and alpha bands represent the carrier frequencies of thalamo-cortical effective connectivity, PSI values were recomputed using a frequency resolution of 2.5-6 Hz and 11-18 Hz for theta and alpha bands, respectively. Finally, as a control measure PSI was additionally computed for the gamma band (30-60 Hz).

#### Spike-spike correlation

Spike cross correlation analysis was used to determine the temporal dependence of co-recorded spiking activity in LP/Pulvinar and PPC. Spike time series in thalamus and cortex were binarized at a sample rate of 1 kHz. Cross correlation functions were then computed on binarized time series for all possible combinations of LP/Pulvinar and PPC channel pairs. To determine the oscillatory structure of spike cross correlations, an FFT was computed on spike cross correlation functions from -0.5 to 0.5 seconds, with oscillatory power defined as the absolute value of the square of the FFT at each carrier frequency. To determine the dependence of synchronized oscillations in spiking on pupil diameter, spike cross correlation functions were computed for spikes occurring during each pupil diameter bin. As above, the oscillatory power of pupil diameter dependent spike correlations was computed using a FFT for all pupil bins.

#### Statistics

All statistical tests were performed in Matlab (Mathworks). To test if neurophysiological and functional connectivity metrics significantly vary with fluctuations in pupil diameter, we computed a one-way ANOVA across the eight pupil diameter bins. For frequency resolved analyses, one-way ANOVAs were computed across the entire frequency spectrum, with p-values corrected for multiple comparisons using false discovery rate (Matlab function *mafdr.m*, with ‘*BHFDR*’ set to ‘*true*’). To quantify the linear relationship between neural dynamics and changes in pupil diameter, we computed the Pearson correlation of neuronal and functional connectivity variables across the eight pupil diameter bins.

#### Code and data availability

Electrophysiological and pupillometry data, as well as MATLAB code that was used to perform outlined analyses, can be made available from the corresponding author upon request.

---

**Supplementary Figure 1 /.**
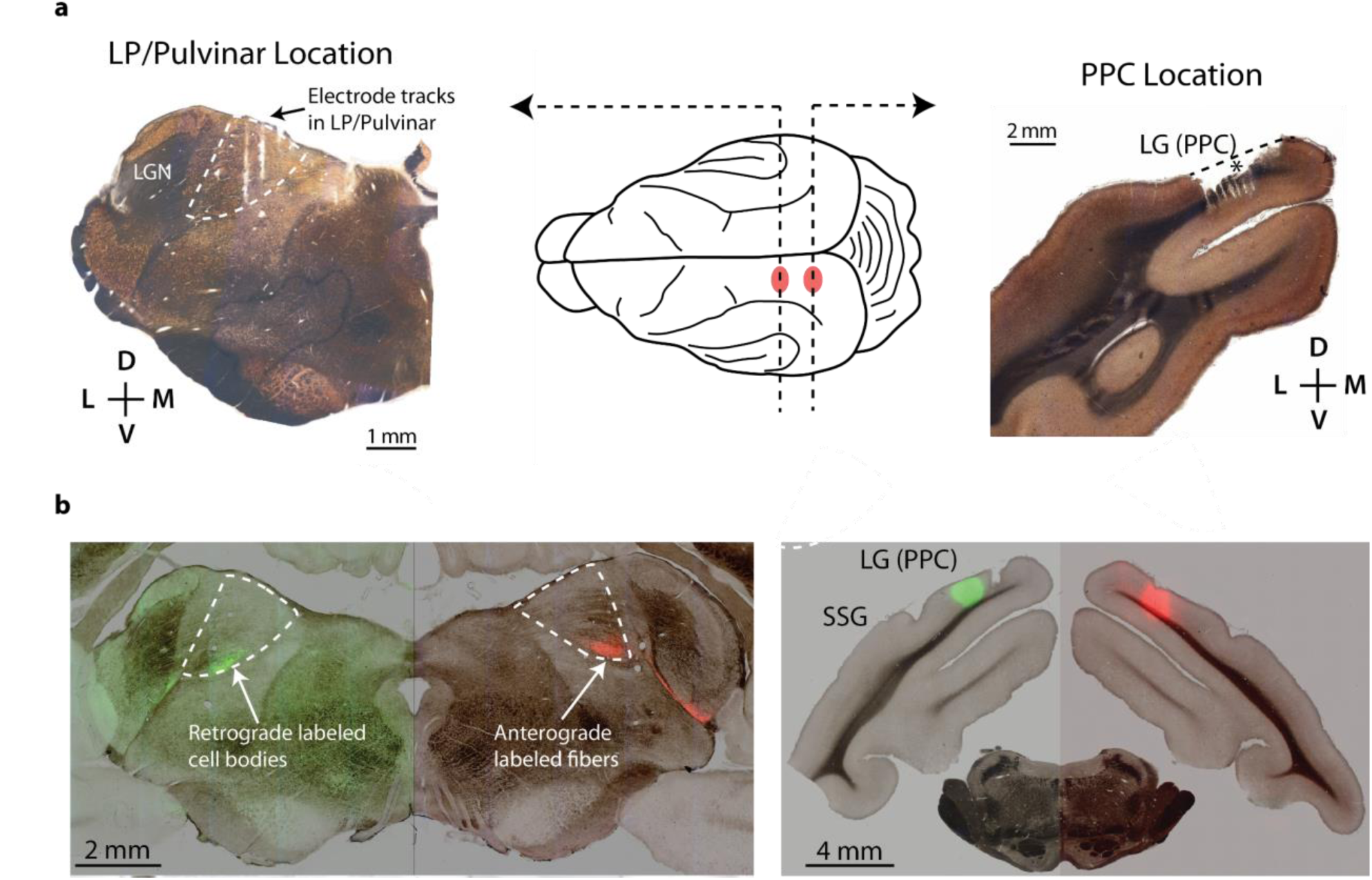
Histological confirmation of multielectrode array recording sites in anatomically connected sites of PPC and LP/Pulvinar. **a**, Coronal sections of LP/Pulvinar (left) and PPC (right) stained for cytochrome oxidase were used for verifying the location of multielectrode array recording sites. The middle panel shows a schematic representation of the ferret brain, with the approximate location of LP/Pulvinar and PPC electrode implantation shown in red. Left: A representative section showing electrode tracks and recording site in the LP/Pulvinar. Right: A representative section showing the recording site of a PPC microelectrode array. The location of the electrode implantation is illustrated by the damage induced from removing the microelectrode array (indicated by *). **b**, The location of LP/Pulvinar and PPC microelectrode implantation overlapped with patterns of anatomical connectivity outlined in previous tracing experiments, indicating that electrophysiological data were obtained from reciprocally connected regions of thalamus and cortex. Abbreviations: LG (lateral gyrus), SSG (suprasylvian gyrus).

**Supplementary Figure 2 /.**
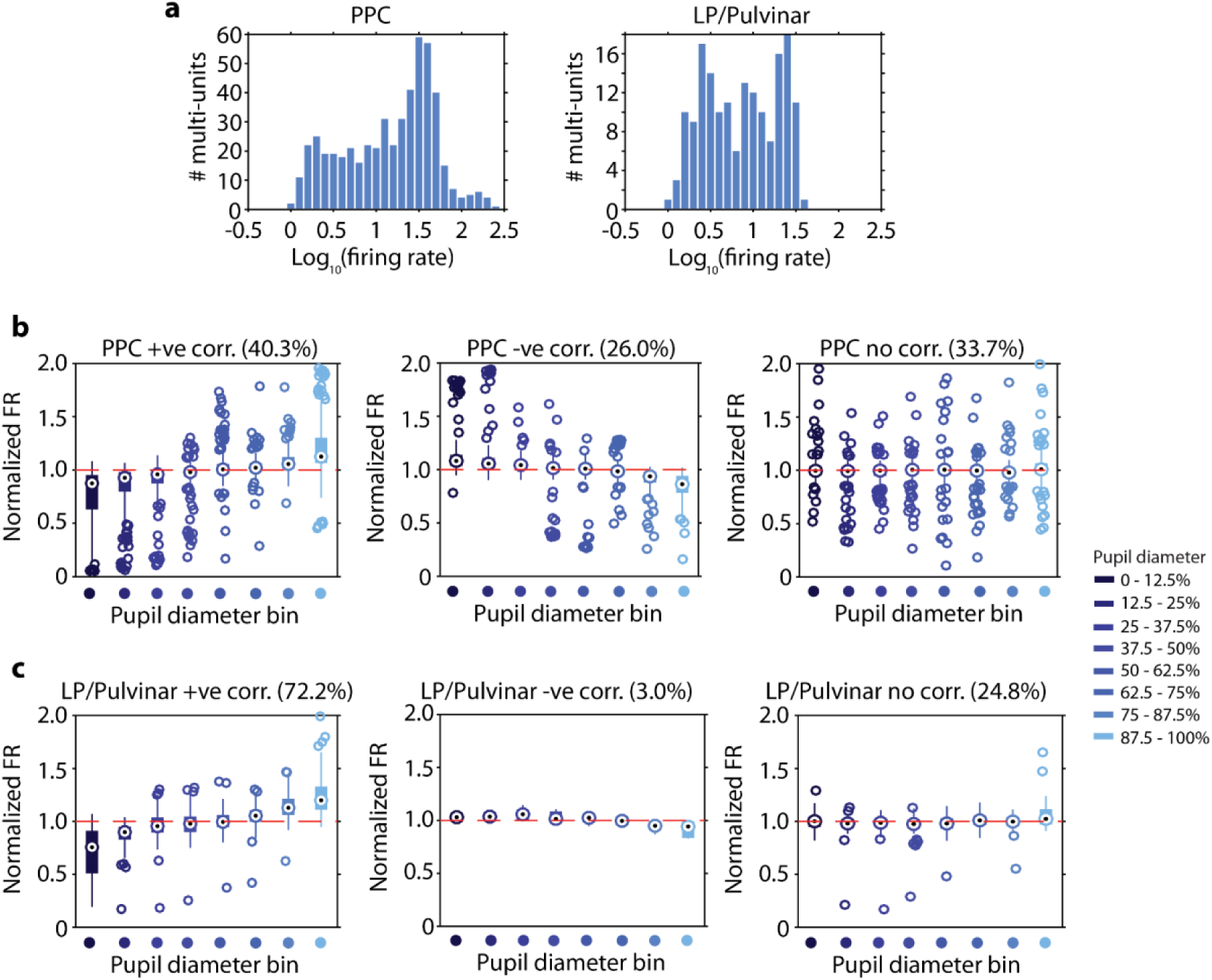
Pupil-linked arousal dependent changes in PPC and LP/Pulvinar firing rate. **a,** Distribution of multi-unit firing rates in PPC (n = 519) and LP/Pulvinar (n = 169). **b**, Population box plots of PPC firing rate across each pupil diameter bin. PPC multi-units were sorted into three populations; units that significantly increased firing rate with increasing pupil diameter (40.3% of units), units that significantly decreased firing rate with increasing pupil diameter (26.0% of units), and units where there was no relationship between firing rate and pupil diameter (33.7% of units). **c**, The same analysis as **b** performed for LP/Pulvinar multi-unit spiking activity. In contrast to PPC, where units displayed both increases and decreases in firing rate with pupil-linked arousal, the vast majority of multi units in LP/Pulvinar (72.2%) displayed significantly increased firing rate with increasing pupil diameter. In addition, very few (3.0%) units displayed a decrease in firing rate with pupil-linked arousal.

**Supplementary Figure 3 /.**
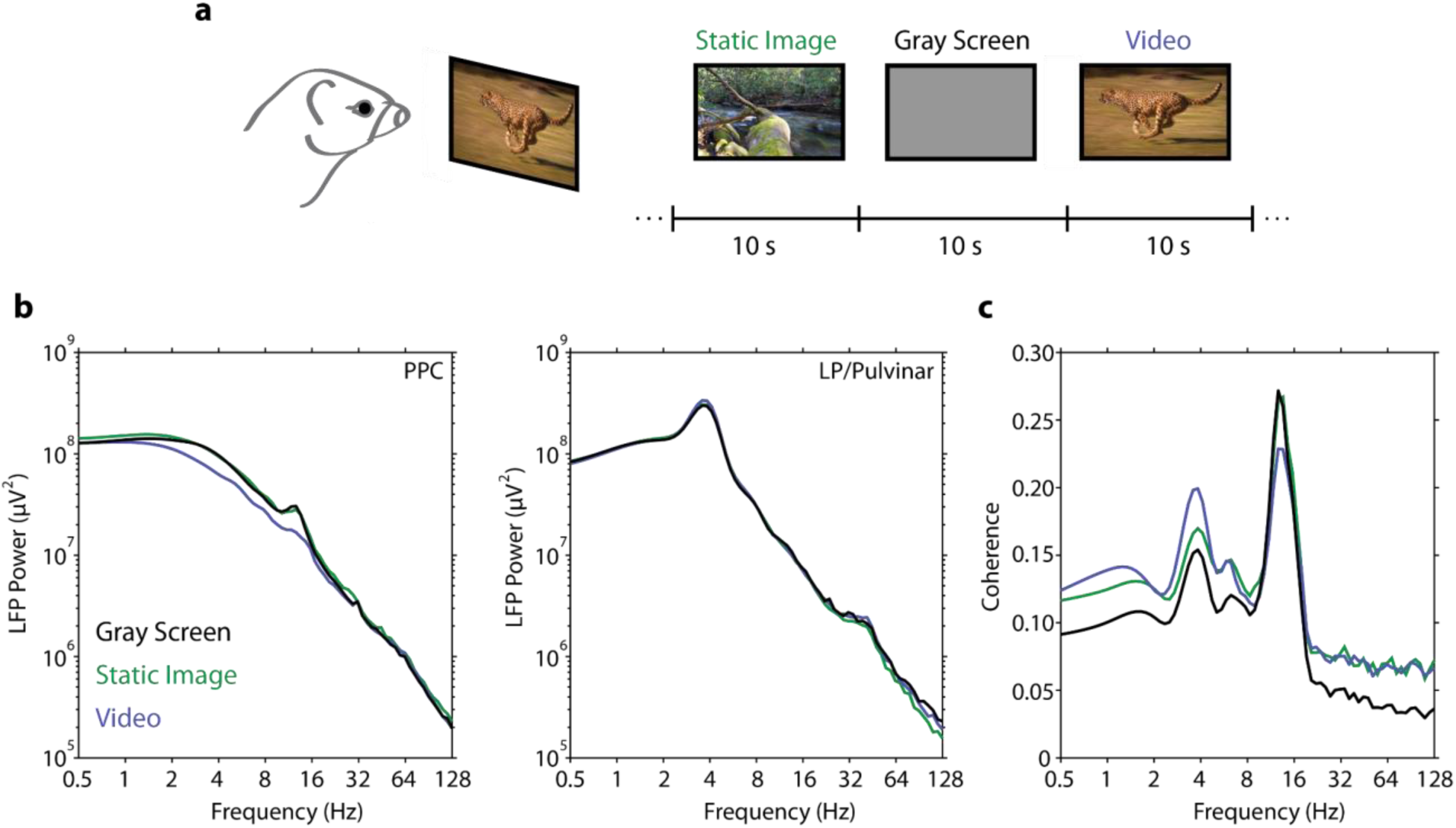
PPC and LP/Pulvinar LFP power spectra under different visual stimulus conditions. **a,** Animals were presented with a library of static images and videos that were randomly interleaved. During the inter-stimulus interval a gray screen was presented. **b**, The mean LFP power spectrum in PPC and LP/Pulvinar for the different stimulus conditions. Note the presence of a prominent peak in the power spectrum at around 12-17 Hz in PPC, which was strongest in the gray screen and static images conditions. LP/Pulvinar LFP spectra were marked by a large peak in the theta frequency band (3.3-4.5 Hz). **c,** The coherence spectrum between co-recorded signals in PPC and LP/Pulvinar. Note the peak in coherence in the 12-17 Hz frequency band that reduces in magnitude for the video condition. Given that LFP signal power and thalamo-cortical coherence in this frequency range was reduced following visual stimulation, we interpreted the frequency band centered around 14Hz to reflect the endogenous alpha frequency in the ferret. This is in line with a mechanistic definition of the alpha rhythm as a thalamo-cortically generated oscillation that reduces in amplitude when networks are engaged in processing sensory information.

**Supplementary Figure 4 /.**
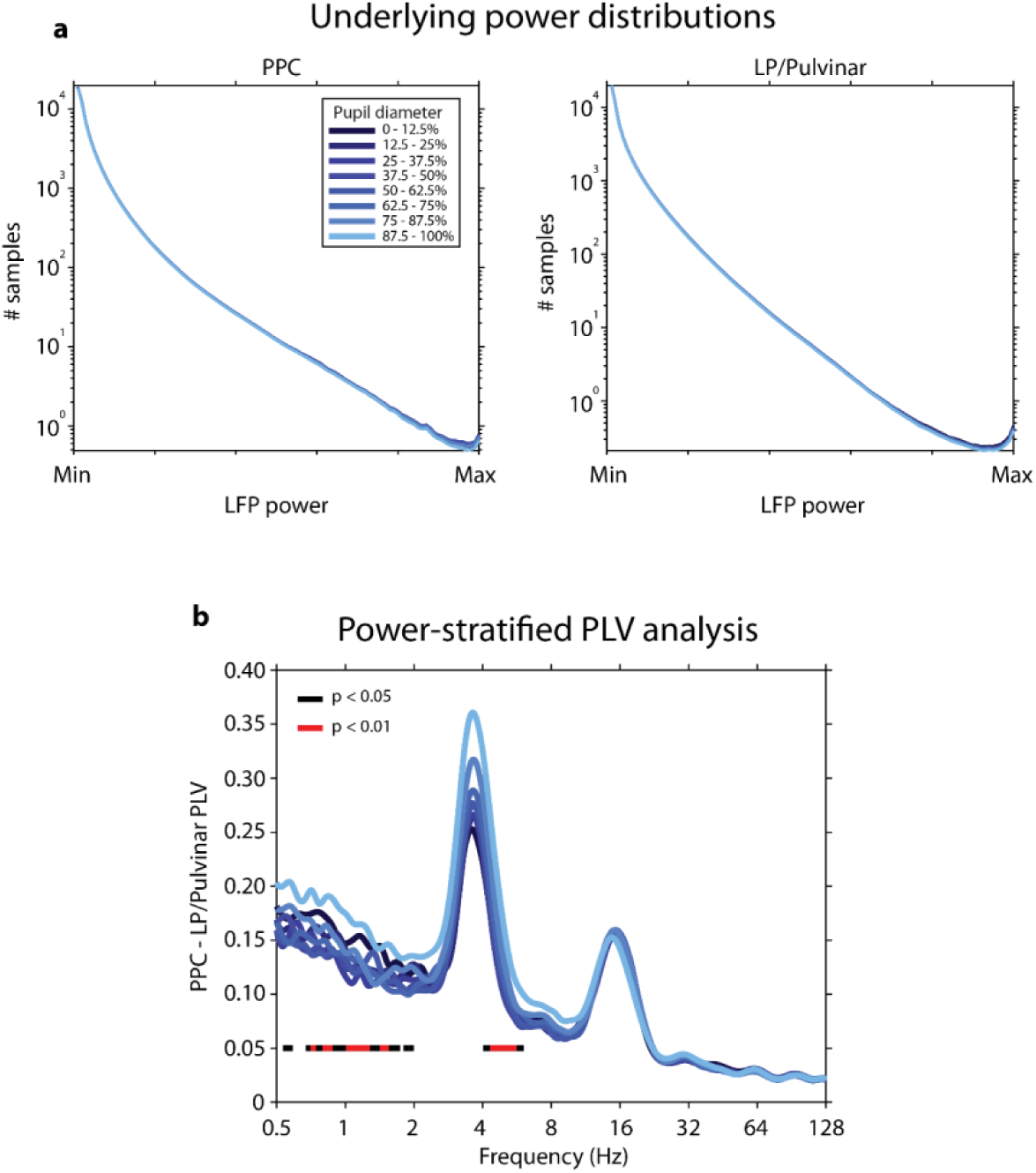
Thalamo-cortical phase synchronization with matched power distributions. **a**, LFP data were subsampled within each frequency band to match the power distributions across all pupil diameter bins. **b**, Thalamo-cortical phase synchronization based on LFP data that were matched for power across pupil diameter bins. The color bars below PLV plots illustrate frequency bands that display a significant effect of pupil diameter (one-way ANOVA, FDR corrected P values). PLV in the theta frequency band remains significant after subsampling data to match power values across conditions. Power matched PLV analysis in the alpha band shows no significant change across pupil diameter bins. However, this result is in contrast to thalamo-cortical spike-spike correlation analysis, which displays a clear effect in the alpha band (Supplementary Figure 5). Given that spike correlations more closely reflect the physiological processes underlying thalamo-cortical functional interaction, we argue that the lack of significant modulation of PLV across pupil diameter bins in the alpha band is an artifact of the power matching procedure.

**Supplementary Figure 5 /.**
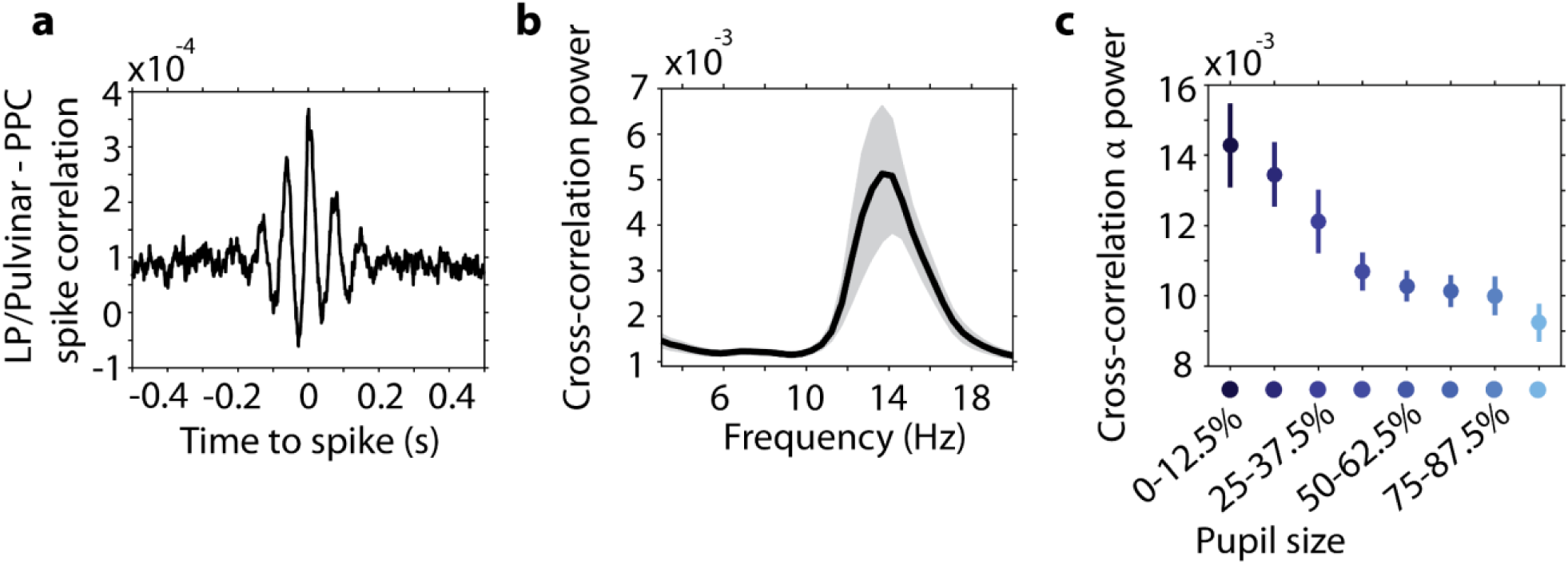
Thalamo-cortical spike correlation as a function of pupil diameter. **a,** Population average spike cross-correlation measured between LP/Pulvinar and PPC recording sites. Note the synchronous oscillatory structure occurring within the alpha frequency band (period of 68 ms ∼14.7 Hz). **b,** The mean power spectrum (± SEM) of thalamo-cortical spike cross-correlation displays a prominent peak in the alpha band. **c,** Alpha oscillatory power in thalamo-cortical spike cross correlation significantly decreased with pupil dilation (P = 4.1^-6^ one-way ANOVA, r = -0.42).

**Supplementary Figure 6 /.**
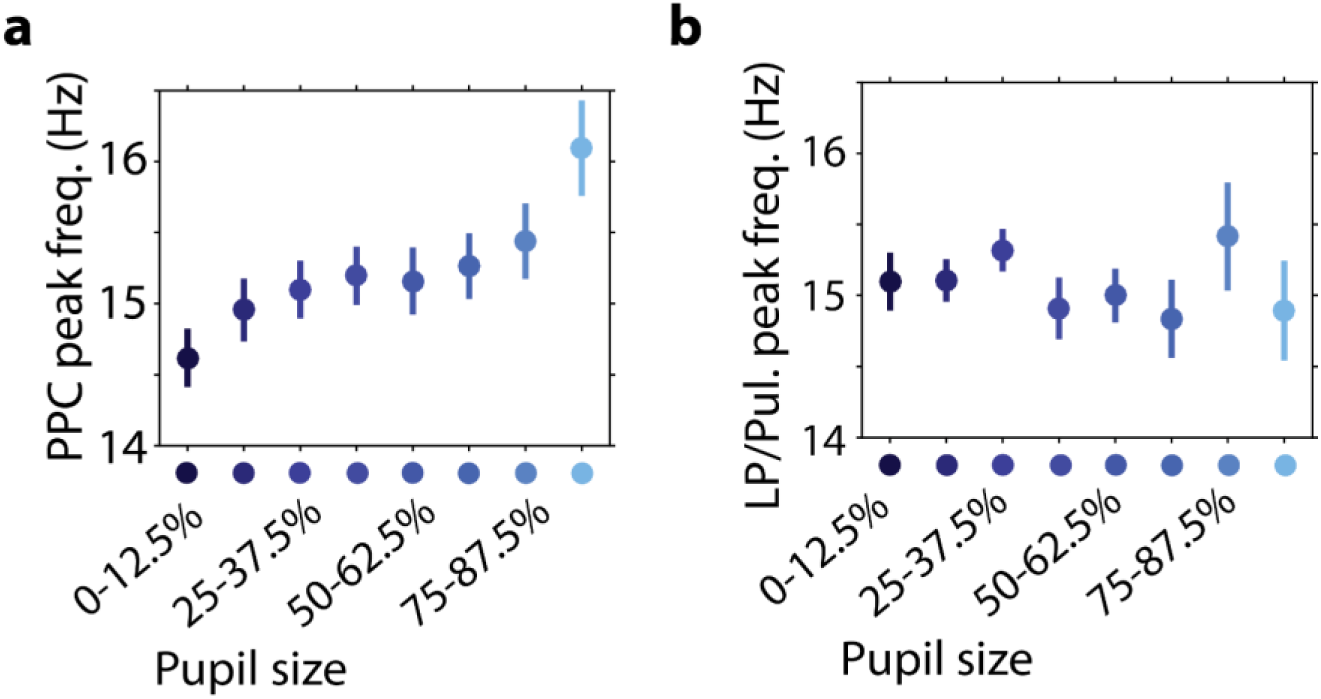
Endogenous alpha frequency speeds up in cortex with arousal. **a-b,** Alpha frequency was measured by taking the peak of spike-LFP phase synchronization spectra within PPC (**a**), and LP/Pulvinar (**b**). Note that peak alpha frequency speeds up in PPC with increasing pupil diameter (P = 0.003 one-way ANOVA). In contrast, peak alpha frequency in LP/Pulvinar displays no relationship to changes in pupil size (P = 0.85 one-way ANOVA).

**Supplementary Figure 7 /.**
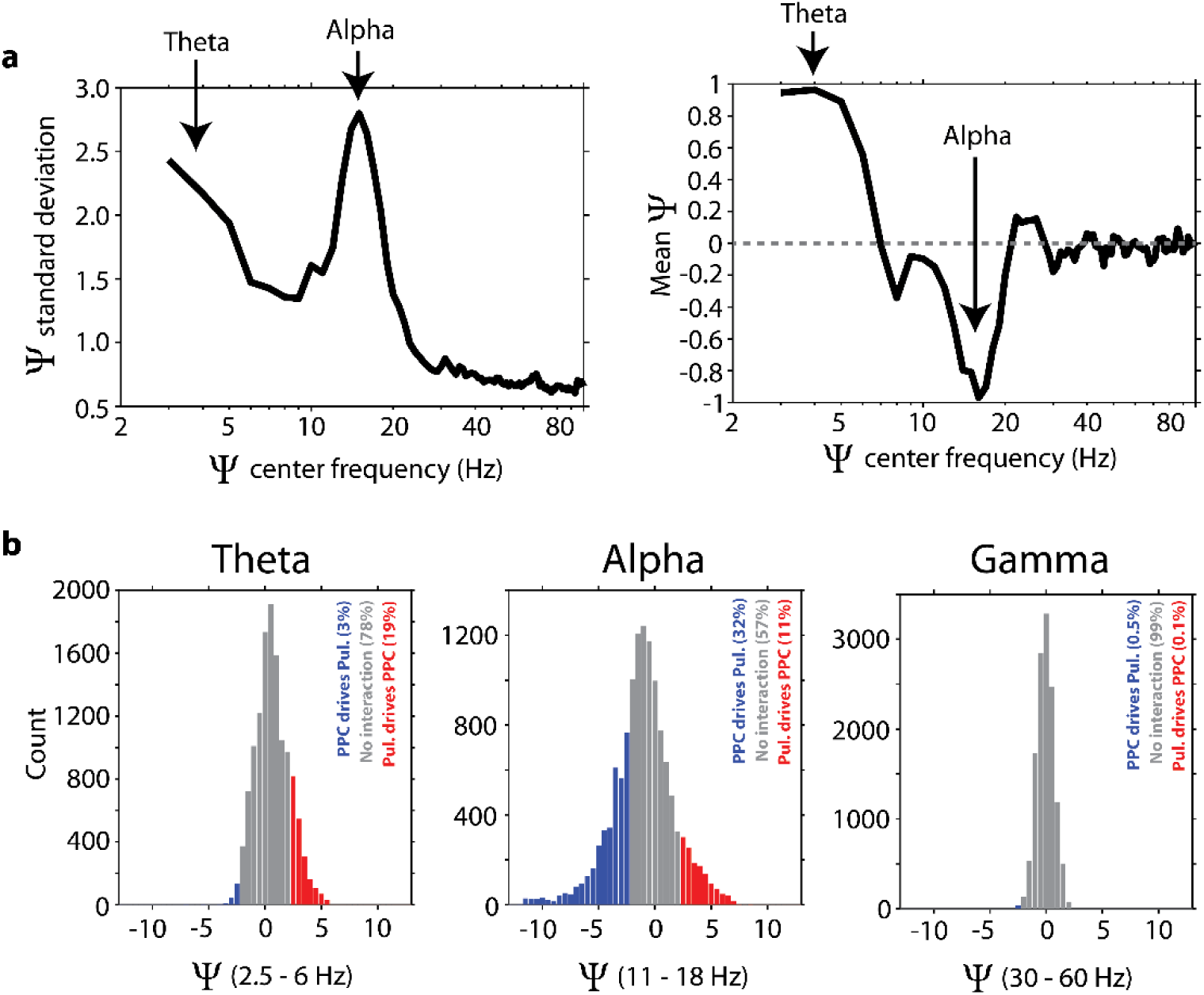
Phase slope index (Ψ) uncovers reciprocal thalamo-cortical effective connectivity in the theta and alpha frequency bands. **a,** The mean and standard deviation of population level phase slope index (PSI) analyses. PSI was initially computed on LP/Pulvinar and PPC LFP signals in a 4 Hz sliding window in the frequency domain (1-100 Hz in 1 Hz steps). We observed both significant drivers and receivers in each recording, with values greater than 2 indicating that LP/Pulvinar significantly drives PPC, and values lower than -2 indicating that PPC significantly drives LP/Pulvinar (p < 0.05). The standard deviation of PSI values was therefore used to infer the width of the PSI distribution, or the extent to which each carrier frequency displays significant drivers or receivers. At the same time the mean indicates if thalamo-cortical (positive PSI values) or cortico-thalamic (negative PSI values) effective connectivity dominates population level PSI analyses. We observed the largest PSI standard deviation in the theta and alpha frequency bands, indicating these are the carrier frequency bands of thalamo-cortical effective connectivity. The mean of the PSI distribution shifted from positive in the theta band to negative in the alpha band, with all other frequencies converging on zero. Together, these results indicate there is generally greater effective connectivity in the theta and alpha bands, but that these frequency bands reflect effective connectivity in opposing directions, with theta band effective connectivity indicating LP/Pulvinar drives PPC, and alpha band effective connectivity indicating PPC drives LP/Pulvinar. **b,** Population level PSI analysis computed on all channel pairs for the theta (2.5-6 Hz), alpha (11-18 Hz), and gamma (30-60 Hz) frequency bands. In each histogram, red bars indicate the proportion of channel pairs where LP/Pulvinar significantly drives PPC, gray bars indicate no significant interaction, and blue bars indicate where PPC significantly drives LP/Pulvinar. The percentage of total channel pairs for each sub-group are given as insets in each figure. Note that theta and alpha PSI values show wider distributions, and that the center of mass is shifted away from zero. In contrast, gamma PSI values are closely centered around zero, indicating this frequency range does not mediate thalamo-cortical effective connectivity.

**Supplementary Figure 8 /.**
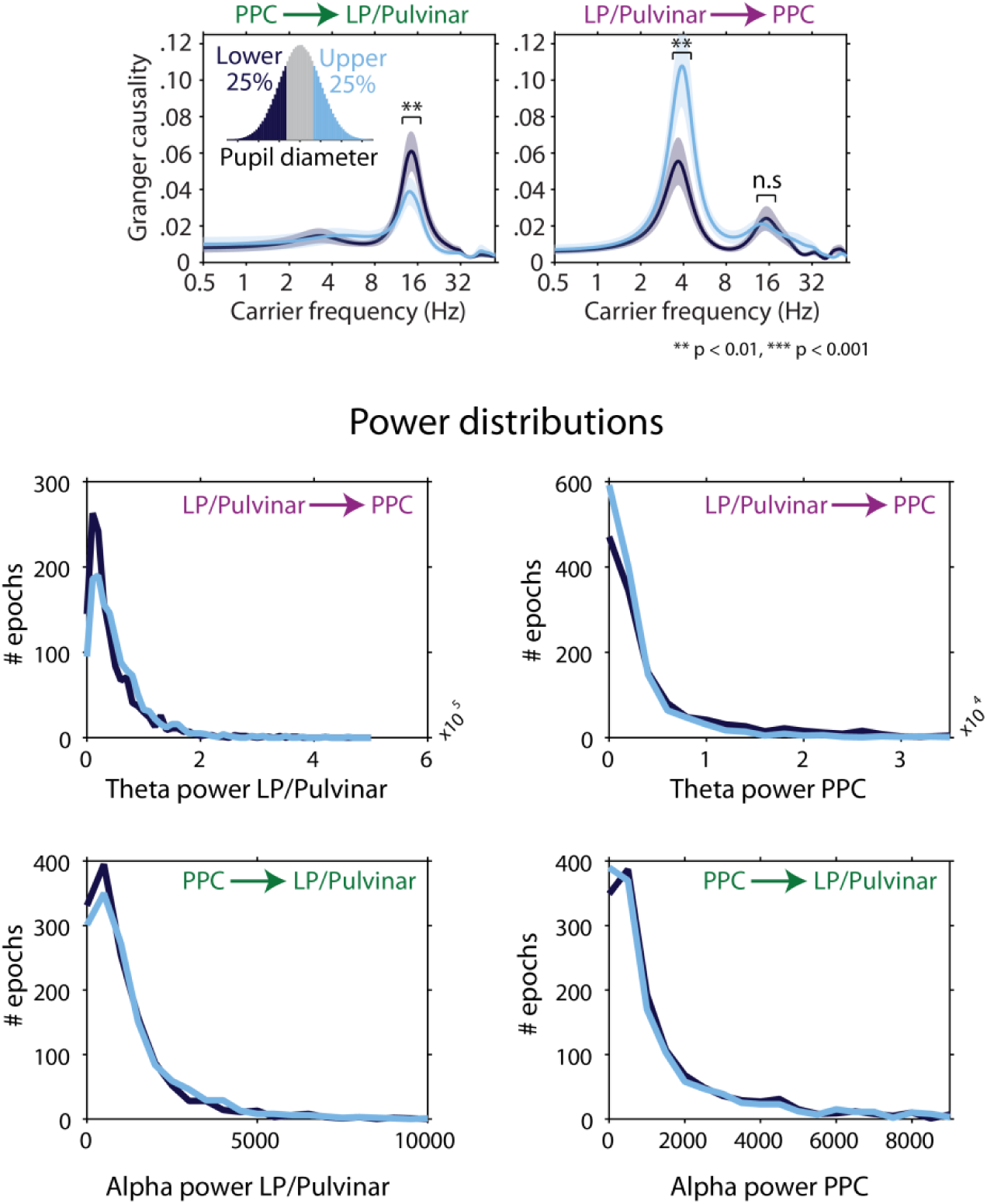
Power matched Granger causality analysis. Granger causality was measured for time periods where the pupil diameter was small (< 25%, dark blue), and large (> 25%, light blue), respectively. Data were subsampled to match power distributions for the theta and alpha frequency bands between conditions. Given that the theta band was the main carrier frequency of thalamus-to-cortex effective connectivity, we matched power distributions in the theta band while computing LP/Pulvinar to PPC Granger causality. Similarly, since alpha was the predominant carrier frequency of cortex-to-thalamus effective connectivity, we matched power distributions in the alpha band while computing PPCto LP/Pulvinar Granger causality. The power distributions of data that were used for Granger causality estimation are shown in the lower four plots.

**Supplementary Figure 9 /.**
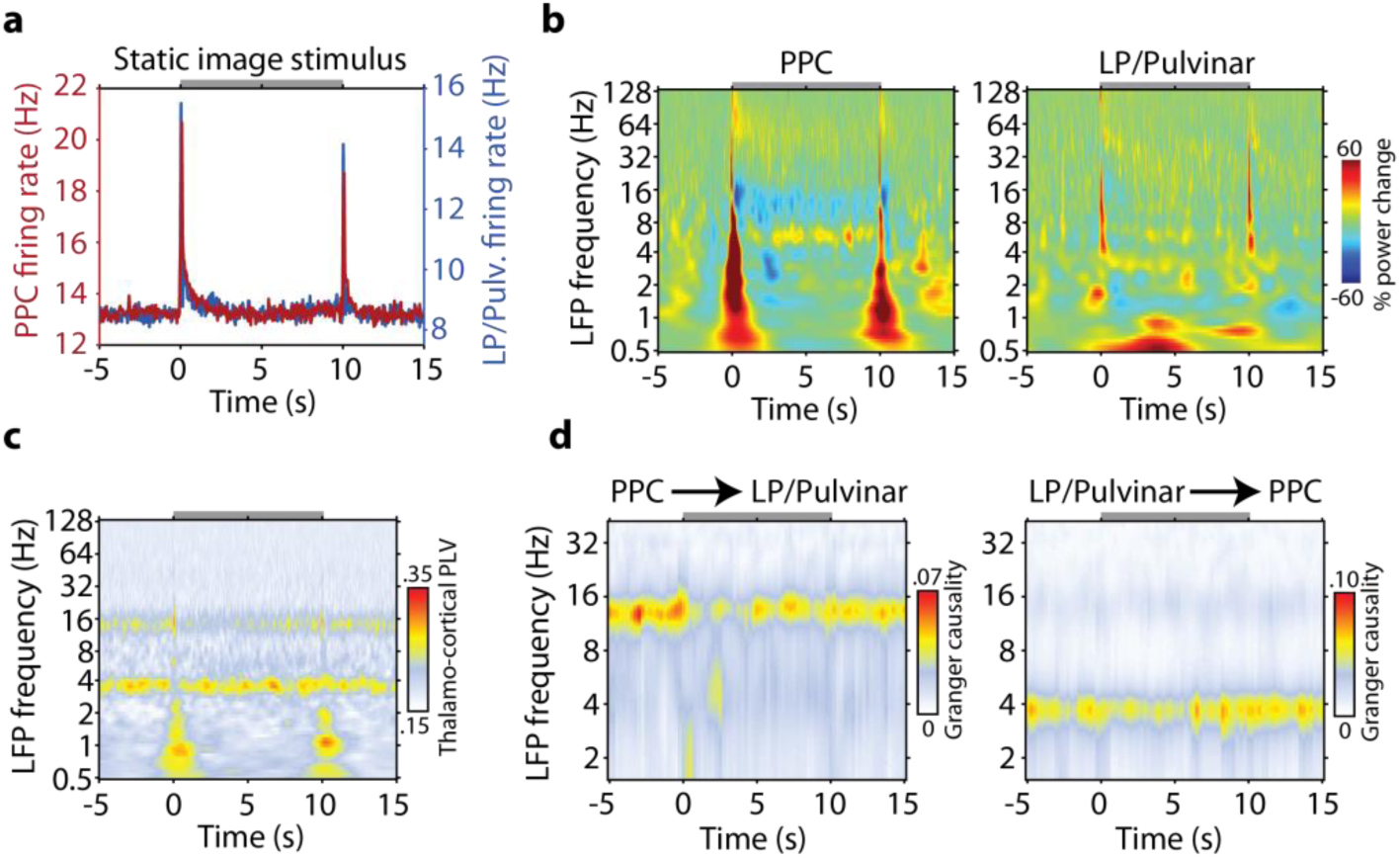
PPC and LP/Pulvinar responses to naturalistic images. **a,** Across session average firing rate during presentation of naturalistic images. **b**, Across session average LFP spectrogram for PPC (left) and LP/Pulvinar (right) in response to naturalistic images. LFP power was normalized by the prestimulus power (-5 to -1 seconds). **c**, Average thalamo-cortical PLV during the time course of naturalistic image presentation. **d**, Average time and spectral resolved Granger causality analysis for naturalistic image presentation. The gray bar at the top of each plot indicates the duration of naturalistic image presentation.

**Supplementary Figure 10 /.**
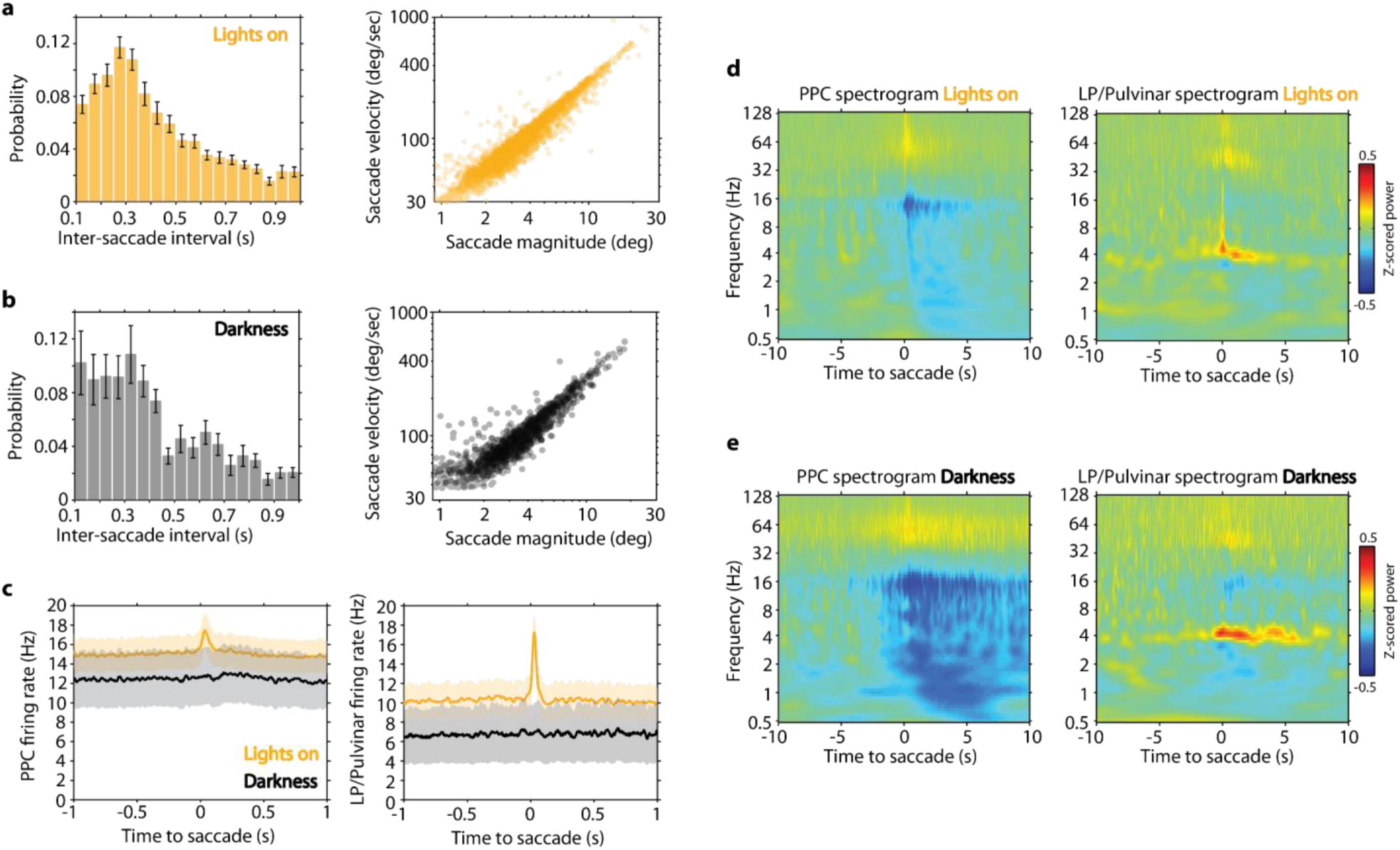
Basic saccade properties and saccade-locked measures of spiking activity and LFP power. **a-b,** Inter saccade interval (± SEM) and scatter plot of saccade magnitude versus peak velocity for saccades performed with the lights on (**a**) and in a dark room (**b**). **c**, Across session average firing rate in PPC (left) and LP/Pulvinar (right) time locked to the occurrence of saccades in the light and dark (shaded regions indicate ± SEM). Saccade-locked increases in firing rate in PPC and LP/Pulvinar are only present when the lights are on, suggesting that these responses depend on visual input. **d-e**, Across session average LFP spectrograms computed around the occurrence of saccades with the lights on (**d**) and in a dark room (**e**). LFP power was z-score normalized within each frequency band across the entire recording session. In contrast to the fast modulation of spiking activity during saccades, LFP power spectra in both lights on and lights off conditions reflect slower fluctuations in PPC alpha and LP/Pulvinar theta rhythms.

**Supplementary Figure 11 /.**
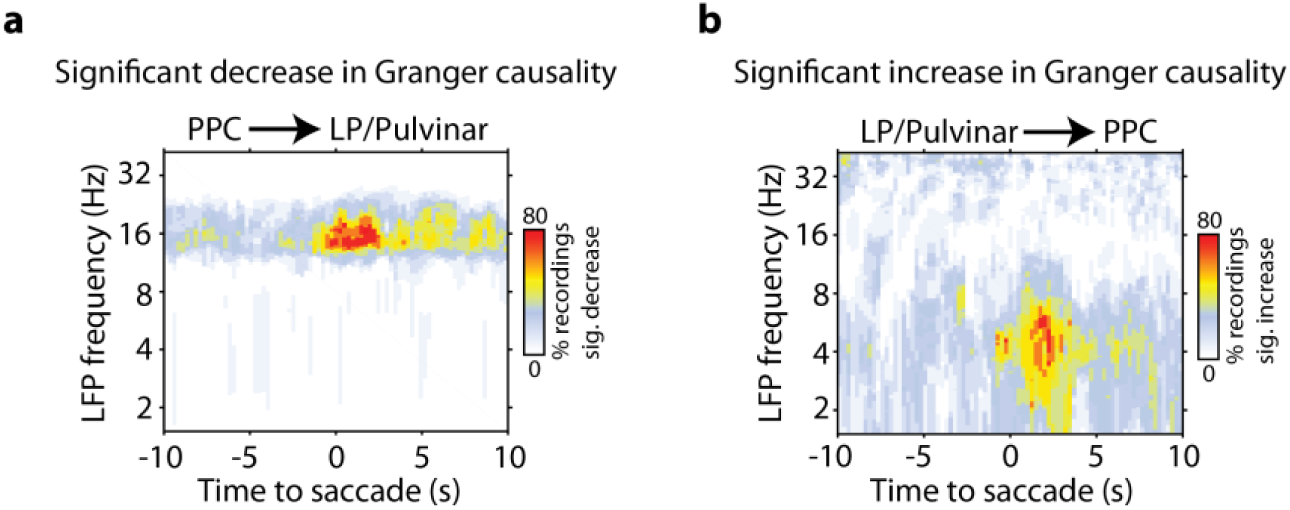
Significant modulation of cortico-thalamic and thalamo-cortical Granger causality time locked to saccades in the dark. **a,** Percent of recording sessions that displayed significant (p < 0.05) reduction in PPC to LP/Pulvinar Granger causality, resolved for frequency and time around the occurrence of saccades. **b**, The same analysis as **a**, but for significant increase in LP/Pulvinar causal influence on PPC. In each case, significance was estimated by computing Granger causality at random time points during each recording, where significant time and frequency points were those that deviated 2 standard deviations from the mean of randomly computed Granger causality estimates. These plots show that the causal influence of cortical alpha rhythms on thalamus breaks down as animals begin to actively sample the visual environment with saccades. Conversely, thalamic causal influence on cortex in the theta band increases predominantly after animals begin to saccade.

**Supplementary Figure 12 /.**
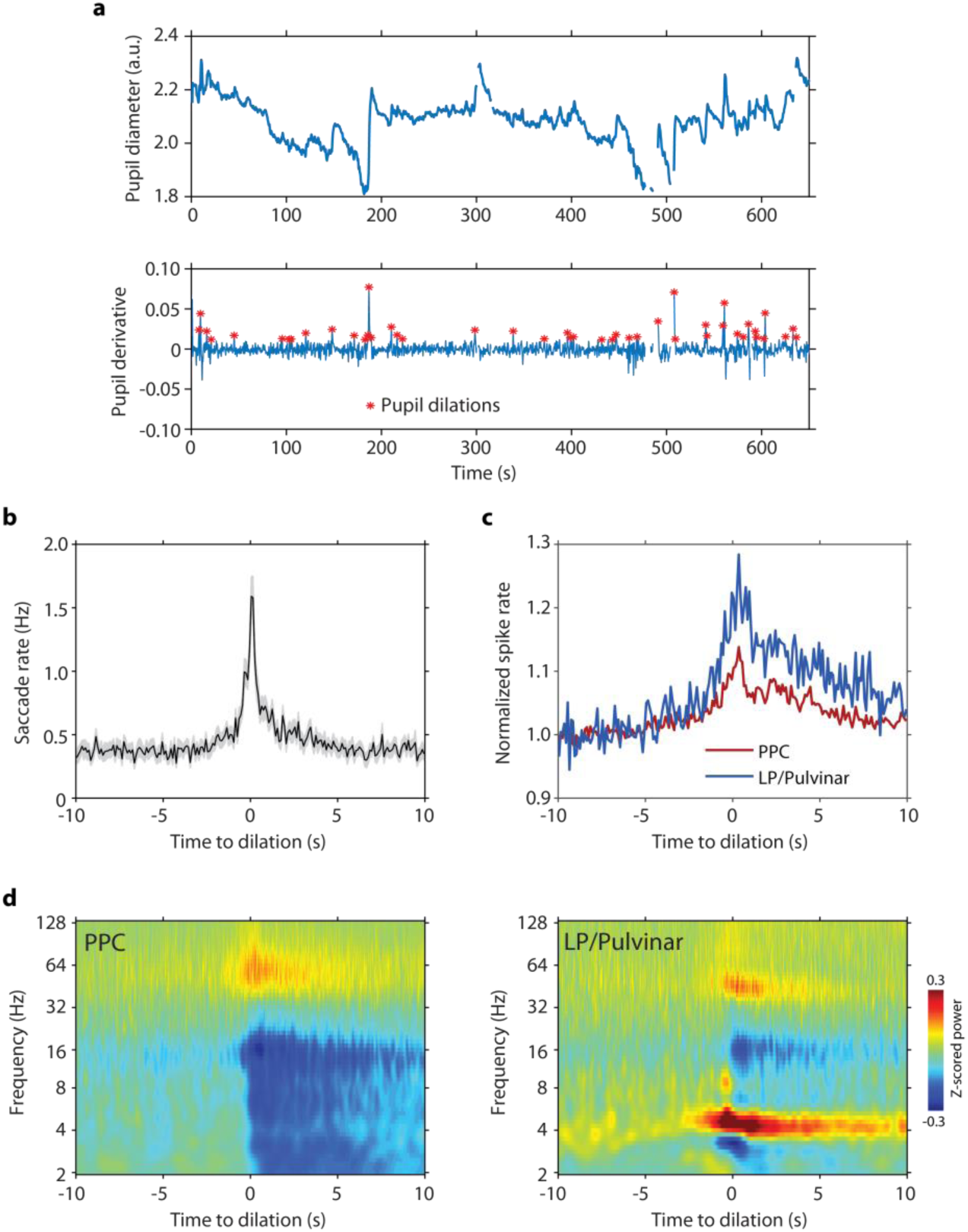
Transient pupil dilations lead to the reorganization of oculomotor behavior and neural activity in PPC and LP/Pulvinar. **a,** Pupil diameter (top) and pupil diameter first derivative (bottom) for one example recording. Transient pupil dilations were detected by finding local peaks in pupil diameter derivative time series. Discontinuities in plots indicate where data have been removed around the occurrence of large-amplitude saccades or eye-blinks. **b**, Mean saccade rate (± SEM) time locked to the occurrence of pupil dilations. **c**, Average multiunit spike rate in PPC and LP/Pulvinar time locked to pupil dilations. **d**, LFP power spectrograms in PPC (left) and LP/Pulvinar (right) time locked to pupil dilations. LFP power was z-score normalized across the entire recording session.

**Supplementary Figure 13 /.**
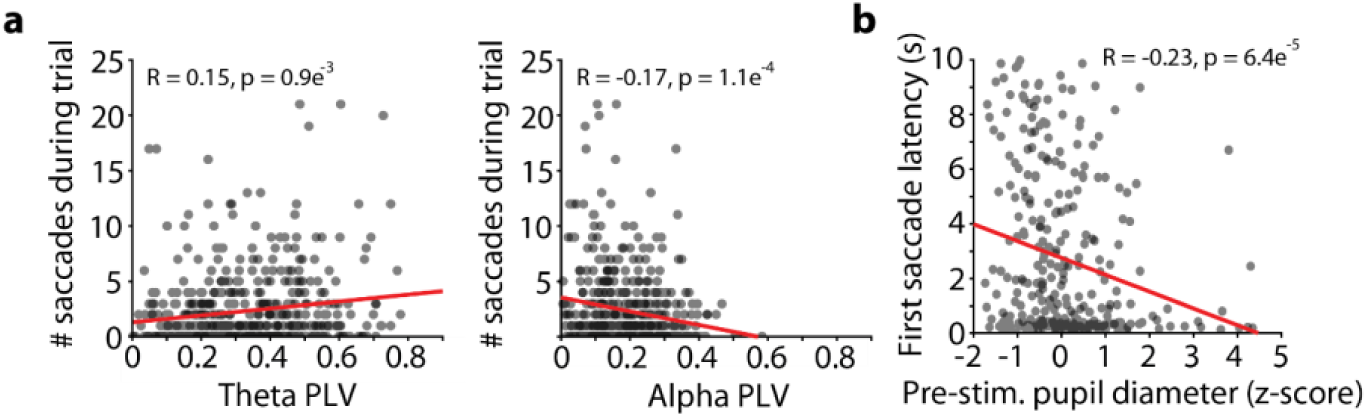
Correlation of thalamo-cortical functional connectivity and prestimulus pupil diameter to oculomotor behavior while viewing naturalistic images. **a,** Correlation of thalamo-cortical phase synchronization in the theta (left) and alpha (right) carrier frequency bands and the number of saccades performed during presentation of naturalistic images. Theta PLV displays a significant positive correlation with saccadic sampling of stimuli, whereas alpha PLV displays a significant negative correlation. **b**, Correlation of pre-stimulus pupil diameter to the latency of the first saccade for subsequent naturalistic image stimulus presentation. Significant negative correlation illustrates that animals sample incoming visual stimuli more rapidly when they are in a more aroused state.

